# Polymicrobial catheter biofilms sustain susceptible *Enterococcus faecalis* and *Escherichia coli* during β-lactam treatment

**DOI:** 10.64898/2026.08.31.748328

**Authors:** Philip A. Karlsson, Meda Skinkytė, Christian Bolin, Huasi Zhou, Himesha Abenayake, Enrique Joffré, Wei Xia, Josef D. Järhult, Helen Wang

## Abstract

Broad-spectrum β-lactam exposure can select for *Enterococcus*-dominated urinary communities in catheterized intensive-care patients, even when co-colonizing *Escherichia coli* remains susceptible. We investigated paired *E. faecalis* and *E. coli* isolates recovered before and after piperacillin-tazobactam (TZP) treatment using a catheter biofilm model and showed that their survival depends on mutualism and biofilm-dependent persistence. Without antibiotics, *E. faecalis* reduced *E. coli* biofilm formation yet promoted pre-attachment coaggregation and reorganized mixed-biofilm architecture on the catheter. Despite TZP susceptibility and the absence of resistance determinants, catheter-associated biofilms and biofilm-dispersed cells survived concentrations 250- to 1000-fold above their MICs, whereas planktonic cells were eliminated. Survivors retained susceptibility but showed delayed regrowth, consistent with a transient persister-like state. In the post-treatment pair, each species sustained the other during recovery, coinciding with a nonsynonymous substitution in the enterococcal surface adhesin Esp. These findings show that antagonistic and cooperative interactions can coexist within catheter biofilms and enable susceptible polymicrobial communities to withstand β-lactam treatment without β-lactam resistance.

## INTRODUCTION

Catheter-associated urinary tract infections (CAUTIs) remain among the most frequent healthcare-associated infections worldwide and constitute a major driver of antibiotic consumption in intensive care units (ICUs) [1]. Indwelling urinary catheters (IDCs) provide a substrate for microbial adhesion and biofilm formation, creating structured environments in which bacteria might be shielded from treatment and host immune responses [2,3].

We recently demonstrated that antimicrobial use during COVID-19 ICU changed the urinary microbiota of catheterized patients [4]. Broad-spectrum β-lactam therapy favored *Enterococcus* as the dominant genus while suppressing common uropathogens, including *Escherichia* and *Klebsiella*. Enterococci accounted for more than 40% of all isolates, representing a marked overrepresentation compared with pre-pandemic cohorts, but in line with similarly reported settings [5–7]. When antibiotic practices shifted during the third pandemic wave, narrower regimens, or no antibiotic treatment, were associated with a decline in enterococcal prevalence, underscoring the link between antibiotic selection pressure and *Enterococcus* dominance [8].

Despite often being regarded as low-virulence commensals [9], or missed in routine diagnostics [10], *Enterococci* display a remarkable ability to persist during antimicrobial therapy, partly due to antimicrobial resistance (AMR), but also to intrinsic resistance, occasionally referred to as tolerance [11–14]. Prior non-lethal exposure to antibiotics can further stunt downstream growth *in vitro* or induce persisters/viable but not culturable cells (VBNC), complicating diagnostic discovery [15–19].

In our previous study, we observed that *Enterococcus* spp. and antibiotic-susceptible uropathogenic *E. coli* (UPEC) frequently co-occurred, even among patients receiving high-dose piperacillin-tazobactam (TZP) [4]. In fact, we could not identify *E. coli* in the urine of any patient unless it co-colonized with *Enterococcus* or was categorized as multidrug-resistant (MDR). Co-isolation of *Enterococcus* and *E. coli* from the urinary tract is increasingly being reported, emphasizing their importance in UTI establishment and pathogenesis [20,21].

TZP accumulates in urine to concentrations lethal even to many resistant bacteria, and how susceptible organisms survive under such conditions remains unclear [22–24]. Biofilm formation is a well-established contributor to antibiotic tolerance, enabling survival at higher antibiotic concentrations in *in vitro* settings [25,26]. However, existing biofilm models rarely capture the clinical setting and are typically grown on polystyrene or polypropylene plates rather than the medical-grade silicone of urinary catheters. Moreover, antibiotic concentrations used in *in vitro* assays rarely match those found in patient urine.

Biofilms form and mature in response to their physical context and surface, while motility and adhesion kinetics of surface–cell interactions jointly determine how communities evolve [27,28]. Non-motile cells, like *E. faecalis*, rely on Brownian motion and gravitational settling for surface encounter, whereas motile bacteria, like *E. coli*, actively increase surface-collision frequency [29,30]. In catheterized systems, motility-mediated surface attachment becomes increasingly important, as non-motile bacteria must rely on early encounters for attachment [28]. In IDCs, the luminal compartment can additionally retain fluid via capillary forces, possibly allowing for different adhesion kinetics. Despite the frequency of CAUTIs in ICUs, silicone IDCs remain less prone to bacterial attachment than the alternatives [31]. Human urinary proteins eventually layer the material in a preconditioned surface (PCS), which can increase downstream colonization efficiencies of subsequent colonizers. Similar PCSs can be expected between primary and secondary bacterial colonizers during polymicrobial infections [32].

UPEC is a flagellated, rapid colonizer of abiotic surfaces and uses type 1 fimbriae (*fim* operon) for initial attachment. Mechanotransduction via irreversible binding causes downregulation of motility genes (including the flagella) and upregulation of general attachment genes, such as curli fimbriae (*csg* operon), P-pilus, and F-pilus. Autotransporters (e.g., Ag43, AidA, and TibA) promote cell-cell aggregation (which does not happen in the planktonic state) and secretion of extracellular polymeric substances (EPS). In *E. coli*, biofilm is often multilayered and mainly consists of exopolysaccharides cellulose (*bsc* operon) and poly-beta-1,6-N-acetyl-D-glucosamine (PGA) (*pga*/*ycd* operon), strengthened by the expression of a negatively charged capsule composed mainly of colanic acid (*wca* cluster) [33,34].

Due to immobility, *E. faecalis* heavily relies on surface adhesins, including the enterococcal surface protein (Esp), adhesin to collagen from *E. faecalis* (Ace), aggregation substance (Agg or Asc10), and possibly biofilm-associated glycolipid synthesis A (BsgA) [35,36]. It has also been shown that the endocarditis- and biofilm-associated pilus (Ebp) (*ebp* operon), is required for binding to fibrinogen-coated catheters [37]. Aggregation substance induces enterococcal clumping, which has been suggested to increase the rate of *E. faecalis* biofilm formation [38,39]. Its biofilm primarily consists of extracellular DNA (eDNA) that surrounds the cells in a “yarn and sweater” structure. Intriguingly, these “spiderweb”-connected cells remain viable during eDNA secretion in early-stage biofilm development [40]. Enterococcal EPS production is limited. One study identified that, except for proteases (e.g., SprE, AtlA, and GelE), the biofilm primarily consists of exopolysaccharides (e.g., stachyose, mannose, maltose, xylose, and lactose) and fatty acids (e.g., palmitic, oleic, stearic, acetic, and butyric acids) [41].

Current studies of *E. faecalis* and *E. coli* primarily indicate a cooperative partnership, yet little is known about their dynamic within catheter biofilms or under antimicrobial stress. It is notable that studies commonly report *E. coli* benefiting from co-growth, whereas other Gram-negative bacteria frequently found in biofilm with *Enterococcus*, including *K. pneumoniae* and *Pseudomonas aeruginosa*, appear inhibited by *E. faecalis* lactic acid production [42,43]. It therefore remains unresolved whether *E. faecalis* assists or antagonizes *E. coli* as the two establish a biofilm on a catheter surface, why and how the two species are consistently recovered together, and whether the resulting community can shelter otherwise susceptible cells from the β-lactam concentrations reached in urine.

Based on our observations and previous literature, we hypothesized that (i) *E. faecalis* aids in the establishment of *E. coli* biofilms on urinary catheters through interspecies interactions and (ii) that catheter-associated polymicrobial biofilms represent the primary protective compartment enabling community-level tolerance against clinically relevant β-lactam concentrations.

To examine these questions, we focused on a prominent case of co-colonization from an ICU patient, HWP028. *E. faecalis* and *E. coli* were co-isolated on the same day before TZP treatment, and again after an 11-day culture-negative interval, providing paired isolates of each species from before and after antibiotic exposure. We employed an in-house catheter-based biofilm assay that preserved luminal geometry and surface characteristics. By integrating this model with clinical context and strain-level phenotyping, the present study aims to elucidate how *E. coli*-*E. faecalis* interactions and biofilm dynamics contribute to bacterial persistence and recurrence in the treated catheterized urinary tract.

## MATERIAL AND METHODS

### Strains, media, and cultivation

Clinical bacterial strains and treatment data used in this study originated from the PronMed COVID-19 intensive care cohort at Uppsala University Hospital [4]. Two clinical strains were used: *E. faecalis* (sequence type (ST) 16, clonal complex 58) and *E. coli* (UPEC O4:H1 ST10309) from patient HWP028, collected longitudinally at multiple time points [44]. These isolates were obtained from the same isolation days, which coincided with collection days nine (HWP028:9) and 14 (HWP028:14) of ICU care. All isolates were routinely maintained on Brain Heart Infusion (BHI) agar (Oxoid), and in BHI broth, incubated at 37°C for 18–24 h under normal conditions. Due to BHI instability at room temperature (RT), which affects *Enterococcus* growth, all BHI was prepared fresh each week, kept completely protected from light, and stored at 4°C for no more than four days, as validated in our lab.

### Whole Genome Sequencing

Fresh bacterial colonies grown on BHI agar were initially identified using matrix-assisted laser desorption ionization time-of-flight mass spectrometry (MALDI-TOF MS) (Bruker Biotyper) and subsequently confirmed by whole-genome sequencing (WGS). Colonies used for MALDI-TOF identification were inoculated into BHI broth and cultured overnight for DNA extraction. Genomic DNA was extracted from overnight cultures using the MasterPure Complete DNA and RNA Purification Kit (Lucigen, Cat. No. MC85200) according to the manufacturer’s instructions. DNA concentration and quality were assessed using a Qubit 2.0 Fluorometer (Thermo Fisher Scientific) with a broad-range double-stranded DNA assay. Extracted DNA was submitted to BMKGENE (Beijing, China) for short-read sequencing using an Illumina-based DNBseq platform with paired-end short-insert libraries (150 bp read length). In addition, long-read sequencing was performed in-house using Oxford Nanopore Technologies with the Rapid Barcoding Kit 24 V14 (SQK-RBK114.24) on an R10.4.1 flow cell using a MinION Mk1D instrument. Base calling was performed using MinKNOW v6.10.1 and Dorado v2.0.1, and long-read quality was assessed using NanoStat v1.5.0. Illumina short reads were quality-filtered and adapter-trimmed using fastp v0.24.0.

### Antimicrobial susceptibility testing

Minimum inhibitory concentration (MIC) was determined by standard broth microdilution according to EUCAST guidelines. Antibiotics were prepared as concentrated stocks in H_2_O (Sigma), sterile-filtered, and diluted to working solutions in cation-adjusted Mueller–Hinton (MH) broth. For TZP, tazobactam was included at a fixed concentration (4 mg/L) while piperacillin was serially diluted. Bacterial inoculum was prepared from fresh 18–24 h colonies, suspended in 0.9% NaCl, adjusted to 0.5 McFarland using a nephelometer, and diluted in MH broth to approximately 10⁶ CFU/mL. Microdilution plates were assembled by dispensing 50 µL MH broth into wells 2–12, adding 100 µL of the antibiotic mixture to well 1, and generating twofold serial dilutions across wells 1–10. Well 11 served as a growth control, and well 12 as a sterility control. Each well received 50 µL of bacterial inoculum (except the sterility control), yielding a total volume of 100 µL. Plates were incubated at 37°C for 16–20 h, and MICs were defined as the antibiotic concentration at which no visible growth was observed. EUCAST-recommended quality-control strains *E. faecalis* ATCC 29212, *E. coli* ATCC 25922, and *E. coli* ATCC 35218 for β-lactam/β-lactamase inhibitor combinations were included, and assays were accepted only when all QC values fell within the EUCAST-defined reference ranges, and the biological replicates did not deviate by more than one dilution.

### Biofilm assays

Biofilm assays were performed using a custom 3D-printed modular tray system adapted from the FlexiPeg [85] and optimized for silicone Foley catheter fragments identical to those used in our ICU cohort (Teleflex Brilliant Plus Aquaflate, 12 Ch, 100% silicone). 3D printing was carried out by U-PRINT, Uppsala University’s 3D printing facility within the Disciplinary Domain of Medicine and Pharmacy. Catheter balloon inflation lumina were sealed with SYLGARD™ 184 silicone elastomer (Dow Corning, 10:1 base-to-curing-agent ratio) to prevent fluid retention in otherwise inaccessible lumina (lumina that cannot be colonized with bacteria in the clinical setting). The catheters were cut into 17 mm segments and mounted vertically in the FlexiPeg-compatible silicone tray, which was aligned with a sterile 96-well microtiter plate. The complete module was placed in autoclave bags, autoclaved at 121°C for 15 min, and dried at 50°C before use. Between experiments, catheters were vortexed in isopropanol, then in 5% Contrad™ 70 detergent solution (Fisher Scientific). Catheters were then rinsed in distilled water until no detergent residues remained and dried at 50°C. Dried pieces were inserted into new modular trays before reuse. Catheters were reused up to eight autoclave cycles as previously internally validated. Any pieces participating in Scanning Electron Microscopy (SEM), showing structural deformation or visible artifacts, were discarded. See the supplementary for step-by-step protocol.

#### Biofilm initiation

Bacterial inoculum was prepared by picking fresh colonies (incubated for up to 24 h), calibrated in 0.9% NaCl using a nephelometer, and diluted to 10⁵ CFU/mL in BHI. Mono-species (*E. faecalis* or *E. coli*) and dual-species (1:1 ratio) biofilms were established in quadruplicates with catheter segments submerged in 160 µL inoculum per well. Plates were incubated at 37°C for 24 h under microaerophilic conditions (CampyGen Compact, Thermo Fisher) inside air-tight containers protected from light. These conditions were intended to better approximate bladder conditions. The catheter module was transferred to a fresh medium or new condition at 24 h unless otherwise stated. At that point, an insert was used to elevate the catheters by 2 mm, and the media volume was adjusted to 180 µL to compensate for the elevation. CampyGen sachets were replaced.

#### Antibiotic exposure and recovery

Piperacillin and tazobactam were dissolved in H_2_O (Sigma), filter-sterilized separately, and stored as concentrated stocks at 40 g/L and 5 g/L, respectively. TZP was mixed into a combined working solution (clinical 8:1 ratio) to yield a final concentration of 4 g/L piperacillin and 0.5 g/L tazobactam. This concentration was selected to model expected urinary TZP levels in ICU patients receiving the standard regimen of 12 g piperacillin and 1.5 g tazobactam per 24 h (**Supplementary Material**) [45–48]. 48 h biofilms (2 x 24 h growth) were transferred into the plate containing TZP and incubated for 24 h at 37°C. After exposure, catheters were transferred to a fresh plate containing antibiotic-free BHI for a 24 h recovery period, and the procedure was repeated for a second consecutive 24 h recovery period (total recovery 48 h), unless otherwise stated. Surrounding media were plated at each time point (post-exposure, after each recovery period). Catheter-associated biofilms were harvested and plated from separate parallel replicate sets corresponding to the time points.

#### Sequential biofilm colonization assays

Catheter segments were first inoculated with a single species (*E. faecalis* or *E. coli*) to allow primary surface colonization. Following 24 h of incubation, catheters were transferred to fresh wells containing BHI inoculated with the second species at standard inoculum. Biofilms were incubated for an additional 24 h, after which the surrounding medium and catheter-associated biofilms were harvested and plated.

#### Conditioned media assay

Conditioned media were prepared by inoculating 30 mL of BHI in a 50 mL conical tube with bacterial suspensions standardized to a 0.5 McFarland standard. Cultures were incubated at 37°C for 24 h with shaking, then centrifuged at 5,000 × g for 10 min at 4 °C to pellet cells and reduce motility. The supernatant was filter-sterilized (0.22 µm). To confirm the absence of viable bacteria, 100 µL of the filtered media was plated onto BHI agar. For biofilm experiments, the conditioned media were used in place of fresh BHI, and biofilms were incubated for 2 x 24 hours. Conditioned media were also used at the 24 h media change.

#### Pre-conditioned surface assay

Catheters in the biofilm tray were submerged in monospecies (*E. coli* or *E. faecalis*) or polymicrobial conditioned media and incubated at 37°C for 24 h to allow potential media compounds to coat the catheter surface. Pre-conditioned catheter segments were transferred into a sterile 96-well plate, each well containing fresh BHI inoculated with *E. coli*. Biofilms were grown for 24 h at 37°C to investigate biofilm initiation.

#### Biofilm growth in human urine and NaCl

BHI was supplemented with pooled human urine at a final concentration of 50%, using urine from recovered patients seen at post-ICU follow-up (group i, see below). After 24 h, catheter segments were transferred to fresh wells containing BHI with 50% urine and incubated for an additional 24 h. To control for the effect of nutrient depletion, parallel experiments were performed with BHI supplemented with 50% 0.9% NaCl in place of urine, following the same protocol.

#### Quantification of viable cells/colony-forming units (CFU)

Catheters were washed through the lumen three times with 200 µL sterile phosphate-buffered saline (PBS) to remove planktonic cells (segments remained submerged throughout), transferred to sterile 2 mL round-bottomed Eppendorf tubes containing 600 µL PBS, and vortexed for 3 min at maximum speed. Ten-fold serial dilutions of each biological replicate were plated in technical quadruplicates on BHI (monocultures) or Brilliance™ UTI Clarity Agar (Oxoid, Thermo Fisher Scientific) to distinguish *E. faecalis* and *E. coli* colonies based on colour differentiation. Incubations occurred overnight at 37 °C.

#### Microscopic imaging through scanning electron microscopy (SEM)

For morphological evaluation, catheters following 48 h incubation were fixed in 2.5 % glutaraldehyde + 1 % paraformaldehyde in Sörensen’s phosphate buffer, dehydrated through graded ethanol (30–100 %), and kept in an airtight box with silica pearls until imaging the same day. Catheter pieces were sputter-coated with Au/Pd (5–6 nm) to improve electrical conductivity for SEM imaging. Secondary electron imaging was performed on a Zeiss Merlin field emission SEM at an accelerating voltage of 3-5 kV, utilizing both InLens and High-Efficiency SE2 (HE-SE2) detectors.

### Aggregation assay

Aggregation was assessed using spectrophotometry, light microscopy, and macroscopic inspection. 1 mL of inoculated broth (10⁵ CFU/mL) containing either *E. coli*, *E. faecalis*, or both species in co-culture was transferred to cuvettes and incubated statically at 37°C for up to 72 h. Optical density at 600 nm (OD600) was measured using a spectrophotometer. Experiments were performed on three independent days, with three biological replicates at each time point. Macroscopic and microscopic assessment of aggregation was performed in parallel. Briefly, 5 mL of inoculated broth was incubated statically in glass tubes at 37°C. At the indicated time points, glass tubes were photographed, and 10 µL samples were collected from the broth surface layer and examined under a light microscope (100x).

### Preparation of urine

For analysis of urine chemistry parameters, equal volumes of urine from ten adult donors were combined to generate representative pools corresponding to four clinical categories: (i) recovered patients seen at post-ICU follow-up, (ii) ICU patients without detectable bacterial growth, (iii) ICU patients with growth of *Enterococcus*, and (iv) ICU patients with non-enterococcal bacterial growth. Pooling and aliquoting were performed under sterile conditions, followed by 0.22 µm filtration before use. Pooled samples underwent standard clinical chemistry analysis for pH, protein, glucose, ketones, leukocytes, nitrite, and blood by Multistix-7, and fibrinogen by immune diffusion anti-fibrinogen antibodies. For experiments, only urine from group (i) was used.

### pH measurements in BHI culture

In vitro pH dynamics were monitored in BHI and in BHI diluted 1:1 with 0.9% NaCl. *E. faecalis*, *E. coli*, and a 1:1 polymicrobial combination of both strains were inoculated at a starting density of 0.5 McFarland units in a total of 15 mL. Uninoculated medium served as a negative control. Cultures were incubated at 37°C in a shaking incubator for up to 48 h, and pH was measured using pH-Fix 2.0–9.0 fixed-indicator strips (MACHEREY-NAGEL, ref. 92118). Each time point was set up as an individual tube, and the experiment was performed in two biological replicates. In addition, the pH of the medium surrounding polymicrobial biofilms was assessed in BHI, in BHI with 1:1 0.9% NaCl, and in BHI with 1:1 pooled human urine, using the same indicator strips, as snapshots at 0 h, 24 h and 48 h. Biofilms were established as described above.

### Investigating persister populations

Regrowth kinetics were measured for *E. coli* and *E. faecalis* HWP028:9 using a Bioscreen C automated growth analyzer (Oy Growth Curves Ab Ltd, Finland). Two cell sources were compared: dispersed cells recovered from unexposed biofilms and dispersed cells recovered from TZP-exposed biofilms (survivors). Biofilms were established and treated as described above. To remove residual antibiotic prior to regrowth, cell suspensions were diluted in cold BHI and centrifuged at 5000 × g for 10 min at 4 °C, and the supernatant was removed, for a total of two washes, followed by resuspension in 1 mL cold BHI. The same washing was applied to every source. Viable counts of each washed undiluted stock were determined by plating serial dilutions on agar.

Each strain was run on a separate 100-well honeycomb plate with BHI at 200 µl per well. To calibrate detection time against inoculum size, the unexposed control was loaded as a tenfold dilution series from 10⁻³ to 10⁻⁸ in four biological replicates. Survivors were loaded undiluted (neat) and at 1:100 in four biological replicates, each with three technical replicates. Sterile BHI wells served as blanks. Plates were kept on ice during loading to synchronize the start of growth, then incubated at 37 °C with continuous shaking, and optical density at 600 nm was recorded every 5 min for 48 h.

For each time point, background was corrected by subtracting the mean optical density of the sterile BHI blank wells. Technical replicates were averaged within each biological replicate, and growth curves were presented as the mean with shaded regions denoting one standard deviation. Growth was calibrated against inoculum size using the unexposed control, with the absolute cell concentration in each dilution well calculated as the viable count of the undiluted stock multiplied by the dilution factor. Time to detection (TTD) was defined as the time at which the background-corrected optical density first reached a threshold of 0.1. TTD was regressed on the base-ten logarithm of the cell concentration, separately for each species, to give the standard curve, with the line of best fit reported as R². The regrowth delay of TZP survivors was quantified against this calibration. For each survivor replicate, the measured time to detection was compared with the value predicted from its viable count, and the difference was taken as the additional regrowth lag. Wells that never reached the detection threshold, and samples from which no colonies were recovered, were excluded.

### Sequence annotation, analysis, and functional genomics

Hybrid assemblies of *E. coli* and *E. faecalis* isolates were generated from Illumina and Nanopore reads using Unicycler v0.5.1 [49] and annotated using Bakta v1.11.4 [50]. Initial pairwise SNP comparisons were performed using Snippy. For higher-resolution comparison of longitudinal isolates, Illumina reads from the later isolate were mapped against the corresponding hybrid assembly of the earlier isolate using minimap2 v2.26 [51] with the short-read preset. Alignments were sorted, indexed, and assessed using SAMtools v1.18. SNPs and small indels were identified using BCFtools v1.18 (mpileup and call) under a haploid model. Candidate variants were filtered using a minimum read depth of 30x, and reference and alternative allele depths were extracted to calculate allele frequencies. Variant coordinates were intersected with the corresponding Bakta GFF3 annotation to classify variants as coding, RNA-associated, or intergenic and to retrieve gene, locus-tag, and product annotations.

Assemblies were uploaded to Proksee (build 2026-02-09) for visualization and comparative genomic analysis. Additional annotations were performed using CARD Resistance Gene Identifier v1.3.1 and AlienHunter v1.3.0 with default parameters. Sequences were additionally analyzed using the Center for Genomic Epidemiology (DTU) web-based platform, including cgMLSTFinder [52,53], ResFinder [54], MobileElementFinder [55], SeroTypeFinder [56], FimTyper [57], and CHTyper [58], to determine strain-specific characteristics. Amino acid sequence alignments were performed using CLC Genomics Workbench v26.0.3 (QIAGEN), and protein domain prediction and functional motif identification were conducted using InterProScan v108.0 (EMBL-EBI) with default settings. Protein structure prediction was performed using the AlphaFold 3 Server (Google DeepMind) with default settings.

### Data processing, statistics, and figure preparation

Data collection, base processing and log₁₀-transformation were performed in Microsoft Excel v16.111.1. Figures and their statistics were produced in GraphPad Prism v11.0.0. Generally, statistical comparisons were performed using unpaired two-tailed t-test with Welch’s correction on log₁₀-transformed CFU values. A significant difference was identified at p-values smaller than 0.05 with * denoting <0.05, **<0.01, ***<0.001, and ****<0.0001. Following initial processing in Excel, pH, growth curves, and the time-to-detection were plotted in Python v3.10.12 using Matplotlib v3.10.9 and NumPy v2.2.6. Plotting code was generated with the assistance of Claude (Anthropic, Opus 4.8) and reviewed and verified by the authors. Figures were assembled in GraphPad Prism, Affinity Designer v1.10.0, and BioRender (2026).

## RESULTS

Paired *E. faecalis* and *E. coli* isolates were obtained by longitudinal urine sampling of a single ICU patient (patient HWP028) with persistent polymicrobial bacteriuria. HWP028:9 and HWP028:14 denote sequential collection indices, not calendar days. HWP028:9 was recovered before TZP treatment. Following TZP, both species became undetectable, and the urine remained culture-negative for 11 days. *E. faecalis* was subsequently recovered alone, followed shortly by the second pair HWP028:14.

### *E. faecalis* impairs *E. coli* biofilm through a soluble factor

In monospecies culture, *E. coli* formed robust biofilms that increased from 24 to 48 h (approx. 800-fold increase for HWP028:9, and 5400-fold increase for HWP028:14, geometric mean). At 24 h, *E. coli* and *E. faecalis* (HWP028:9) reached similar biofilm densities, but only the latter *E. faecalis* isolate (HWP028:14) increased slightly from 24 to 48 h (**Figure 1A**). In dual-species culture, *E. coli* biofilm was unchanged at 24 h but reduced at 48 h relative to monoculture most markedly for HWP028:14 (approx. 2-fold for HWP028:9, and 40-fold for HWP028:14, geometric mean) (**Figure 1B, C**). *E. faecalis* showed the opposite response, increasing in dual-species biofilms at both 24 and 48 h (**Figure 1A, B**). At 48 h, this increase reached more than 10-fold for HWP028:9 and more than 20-fold for HWP028:14 (**Supplementary Figure 1A**).

**Figure 1.**
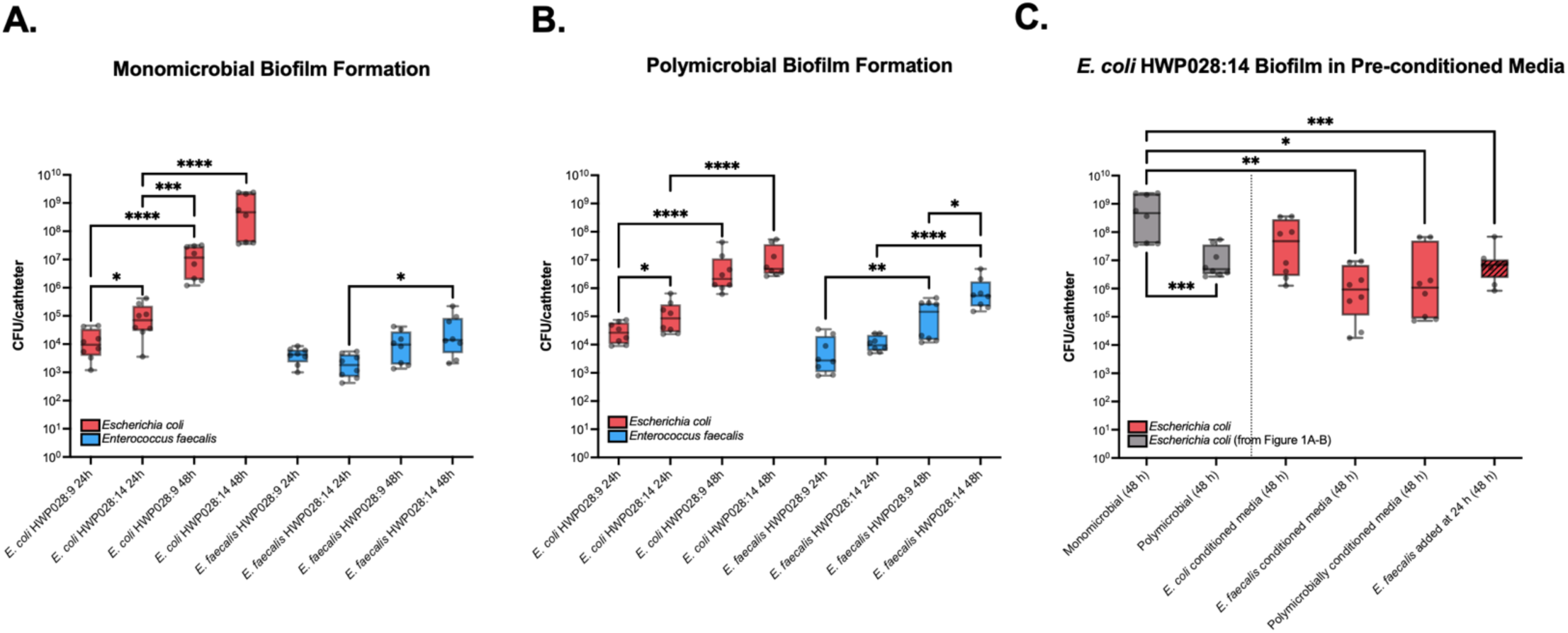
Monospecies and polymicrobial catheter biofilm formation. Biofilm formation was quantified as CFU per catheter after 24 h and 48 h of growth for isolates obtained from two sampling time points (HWP028:9 and HWP028:14). *E. coli* is shown in red and *E. faecalis* in blue. The y-axis displays total CFU per catheter. Box-plot bars show the min-max of eight biological replicates, with the central line representing the median and the error bars indicating SD. Statistical comparisons were performed using an unpaired two-tailed t-test with Welch’s correction on log₁₀-transformed CFU values. **A.** Monospecies growth. **B.** Polymicrobial growth. **C.** *E. coli* HWP028:14 biofilms in pre-conditioned (spent) media and sequential colonization with *E. faecalis* added at 24 h (dashed). Data from Figure 1A-B are reused for comparative reasons (grey).

To identify the basis of the *E. coli* biofilm reduction seen in co-culture, we tested spent (conditioned) medium and sequential inoculation using HWP028:14, the pair with the largest inhibition. *E. faecalis* spent medium reproduced the reduction on *E. coli* biofilm, and spent medium from polymicrobial culture did so to a lesser degree (**Figure 1C**). *E. coli* grown in its own spent medium was unaffected. Adding live *E. faecalis* to an established *E. coli* biofilm at 24 h reduced *E. coli* biofilm to a similar extent as *E. faecalis* spent medium (**Figure 1C**). Preconditioning the catheter with *E. faecalis* spent medium and then inoculating *E. coli* in fresh BHI had no significant effect at 24 h (**Supplementary Figure 1B**).

### *E. faecalis* adopts a distinct architecture in the presence of *E. coli*

To resolve the spatial organization of these biofilms, scanning electron microscopy (SEM) was used. In monoculture, *E. faecalis* HWP028:9 colonized the catheter surface as sparse, discrete microcolonies of typically 100 to 300 cells (**Figure 2A, B**). The microcolonies were separated by tens to hundreds of micrometres, and the cells showed the characteristic diplococcal morphology, attaching predominantly as monolayers with little visible EPS. By contrast, *E. coli* HWP028:9 monocultures formed dense, extensive biofilms of thousands of cells (**Figure 2C**). The matrix occasionally obscured the boundary between the silicone substrate and the biofilm. At higher magnification, individual *E. coli* cells were embedded in a rough, channelled EPS that bridged adjacent cells and anchored them to the catheter (**Figure 2D**).

**Figure 2.**
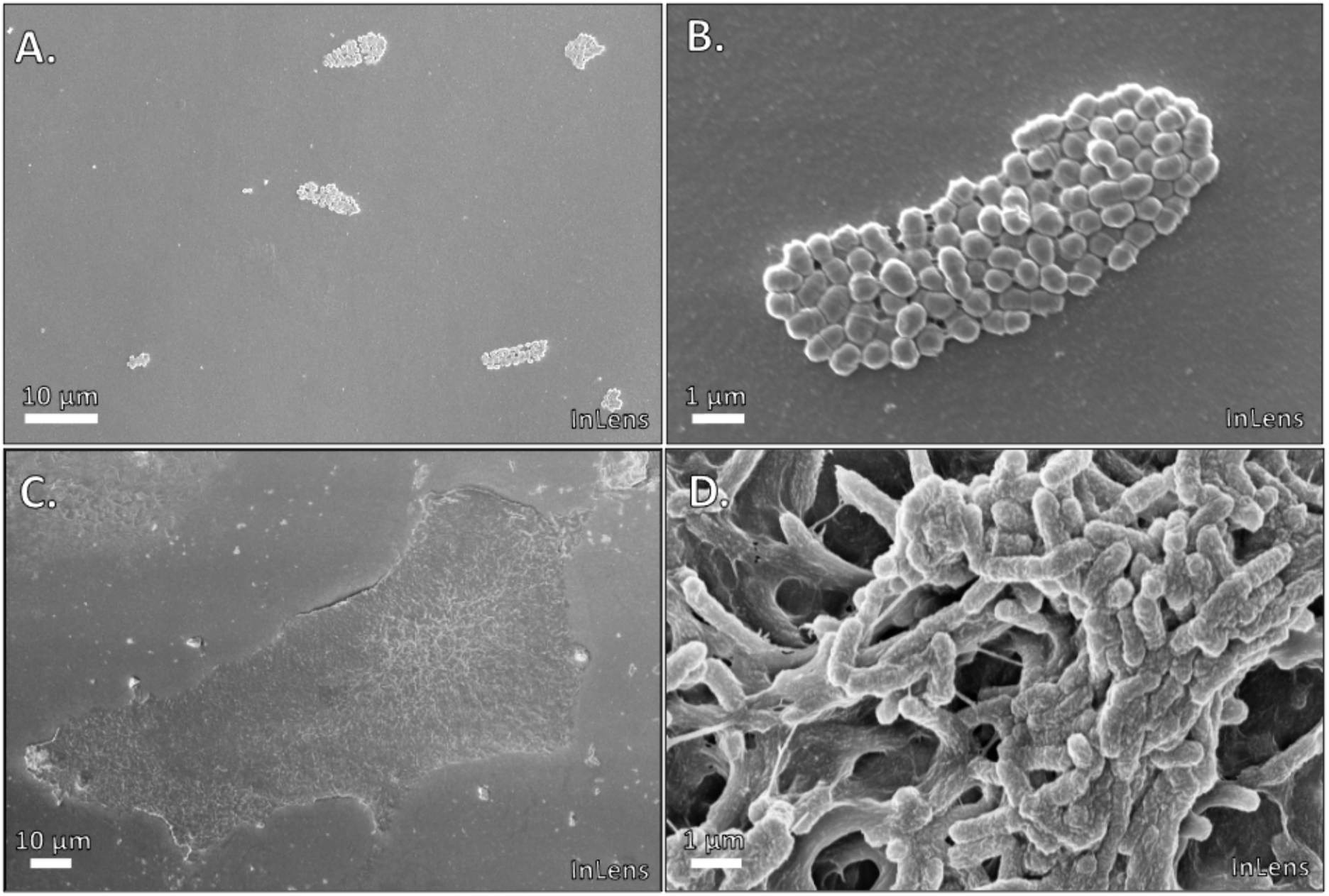
Ultrastructure of monospecies catheter biofilms. Representative scanning electron microscopy of catheters following monospecies 48 h incubation of *E. coli* and *E. faecalis* HWP028:9 in brain heart infusion. **A.** Patch with sporadically attached clusters of *E. faecalis*. **B.** Higher magnification image of *E. faecalis* cluster. **C.** Patch showing layered *E. coli* biofilm. **D.** Higher magnification image of *E. coli* cluster showing multilayered channelling.

Across the polymicrobial catheter surface, SEM revealed marked spatial heterogeneity, comprising four morphologically distinct biofilm types. Two recapitulated the monoculture phenotypes: monoculture *E. faecalis* patches with the cluster phenotype described above, and monoculture *E. coli* patches. The other two were mixed. In *Enterococcus*-dominated regions (**Figure 3A, C, D**), rare *E. coli* rods were interspersed among the enterococcal cells. In *E. coli*-dominated regions (**Figure 3B**), *E. coli* EPS formed the structural scaffold, with *E. faecalis* scattered throughout and embedded in the matrix.

**Figure 3.**
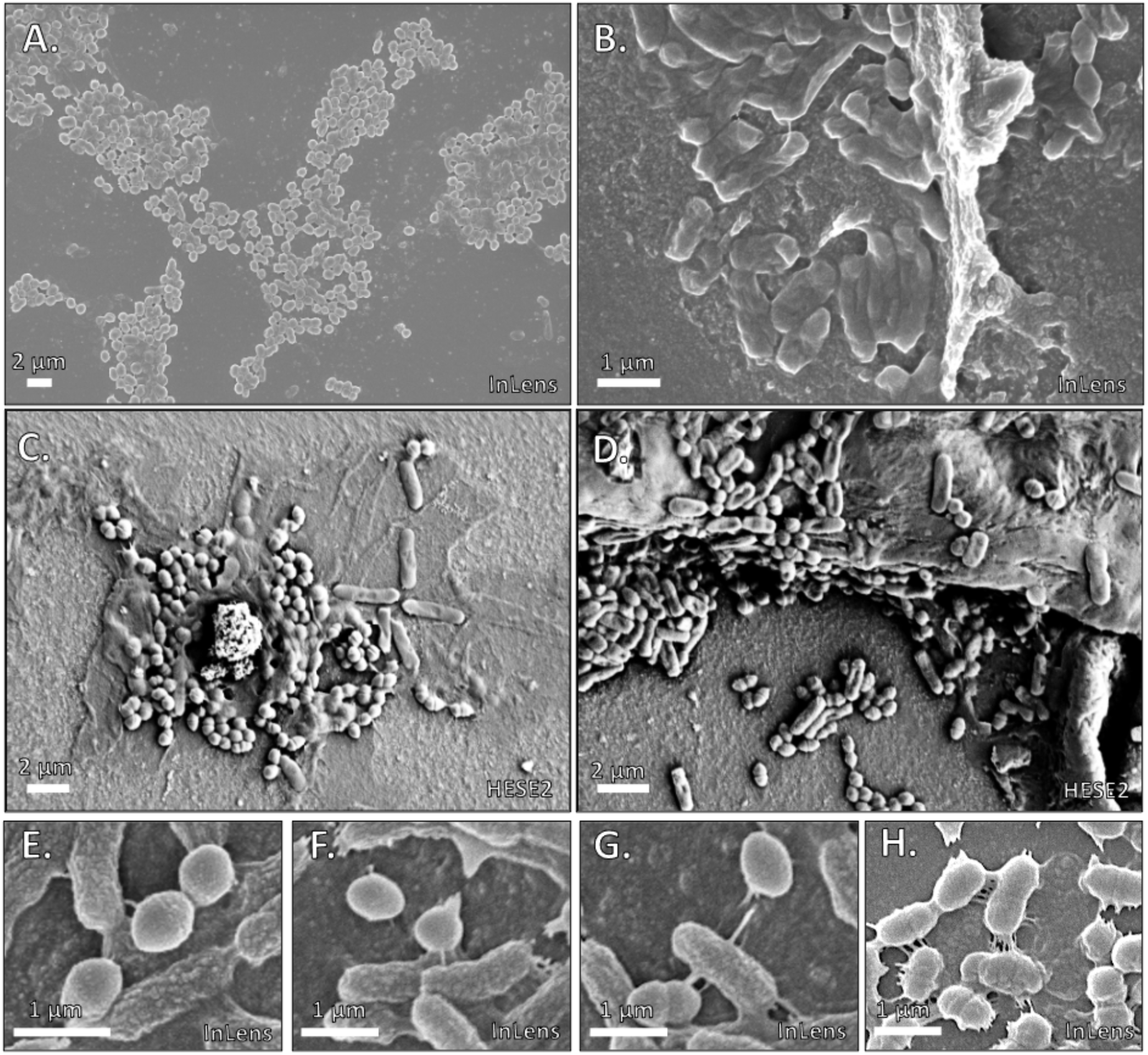
Ultrastructure of polymicrobial catheter biofilm formation. Representative scanning electron microscopy of catheters following polymicrobial 48 h incubation of *E. coli* and *E. faecalis* HWP028:9 in a 1:1 ratio in brain heart infusion. **A.** Patch with visibly spread-out *E. faecalis* captured by InLens detector. **B.** Patch with *E. faecalis* included inside the *E. coli* matrix captured by InLens detector. **C-D.** Examples of *E. faecalis* dominated biofilm captured with High-Efficiency SE2 detector. **E-H.** Example interactions between *E. coli* and *E. faecalis*.

In the *Enterococcus*-dominated regions, *E. faecalis* no longer formed the discrete clusters seen in monoculture but was dispersed as individual cells across the surface, and in places apparently over the *E. coli* matrix, now with visible EPS attaching it to both the catheter and *E. coli* (**Figure 3E-H**). In the *E. coli*-dominated regions, the *E. coli* matrix appeared markedly less developed than in monoculture, and multilayered structures were rare.

### Interspecies co-aggregation occurs independent of surface attachment

The embedding of *E. faecalis* within *E. coli*-dominated regions, and the presence of rare *E. coli* in *Enterococcus*-dominated regions, despite the overall reduction in *E. coli* biofilm, raised the question of whether the two species associate physically, independent of surface attachment. To test this, we monitored aggregation over 72 h in mono- and co-culture (**Figure 4A**). In monoculture, *E. faecalis* auto-aggregated strongly with OD₆₀₀ peaking at approximately 8 h (**Figure 4B**), then declined sharply to near zero by 48 h, consistent with sedimentation, leaving the broth surface largely devoid of cells (**Figure 4H**). *E. coli* monocultures maintained stable high turbidity throughout and remained uniformly dispersed at all time points (**Figure 4C, I**). In co-culture, turbidity dipped slightly below the *E. coli* monoculture between 12 and 20 h (**Figure 4A**), coinciding with the onset of *E. faecalis* sedimentation. Light microscopy at 8 h revealed mixed-species clusters in which *E. coli* rods were embedded within *E. faecalis* aggregates (**Figure 4E–G**). At 48 h, the co-culture surface layer still retained a sparse population of planktonic *E. coli* (**Figure 4J**).

**Figure 4.**
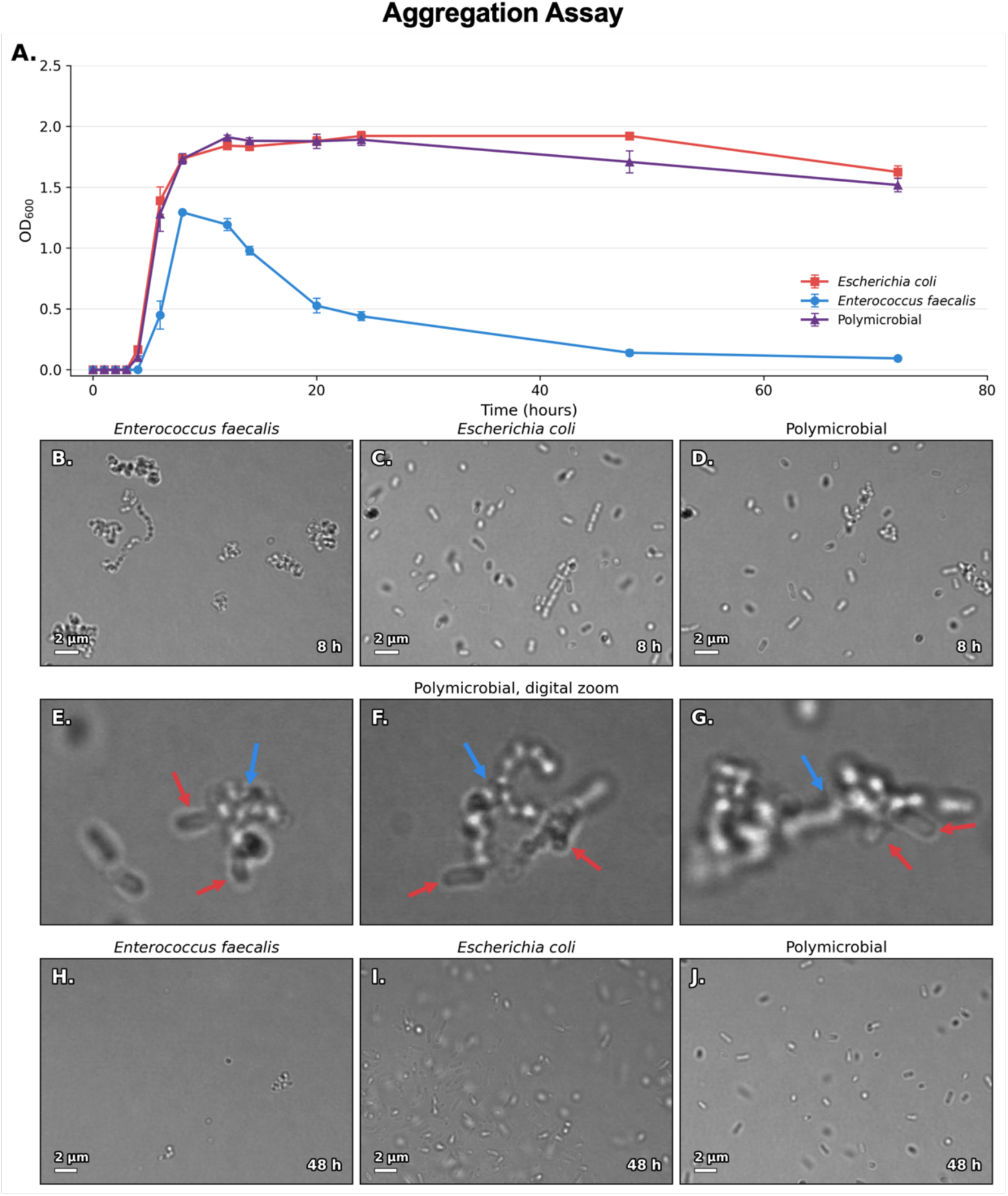
Aggregation behavior of *E. coli* and *E. faecalis* in mono- and co-culture. **A.** Optical density (OD600) measured over 72 h in static monocultures of *E. coli* (red) and *E. faecalis* (blue), and polymicrobial co-culture (purple). Data points represent the mean of nine biological replicates across three independent experiments, and error bars indicate the standard error of the mean. **B–D.** Light microscopy images of the broth surface layer at 8 h from *E. faecalis* monoculture (B), *E. coli* monoculture (C), and polymicrobial co-culture (D). **E–G.** Light microscopy images with digital post-zoom of polymicrobial co-culture samples at 8 h, showing representative mixed-species aggregates. Red arrows indicate *E. coli* rods, blue arrows indicate *E. faecalis* aggregates. **H–J.** Light microscopy images of the broth surface layer at 48 h from *E. faecalis* monoculture (H), *E. coli* monoculture (I), and polymicrobial co-culture (J).

### *E. faecalis* partner determines *E. coli* inhibition in urine

In urine, *E. coli* formed similar monoculture biofilms for HWP028:9 and HWP028:14. Under polymicrobial conditions, *E. coli* HWP028:9 was significantly reduced relative to its monoculture (approx. 800-fold, geometric mean), whereas *E. coli* HWP028:14 was not (**Figure 5A**). *E. faecalis* gained insignificantly from polymicrobial growth in urine, and HWP028:14 grew significantly less than HWP028:9 (**Figure 5A**).

**Figure 5.**
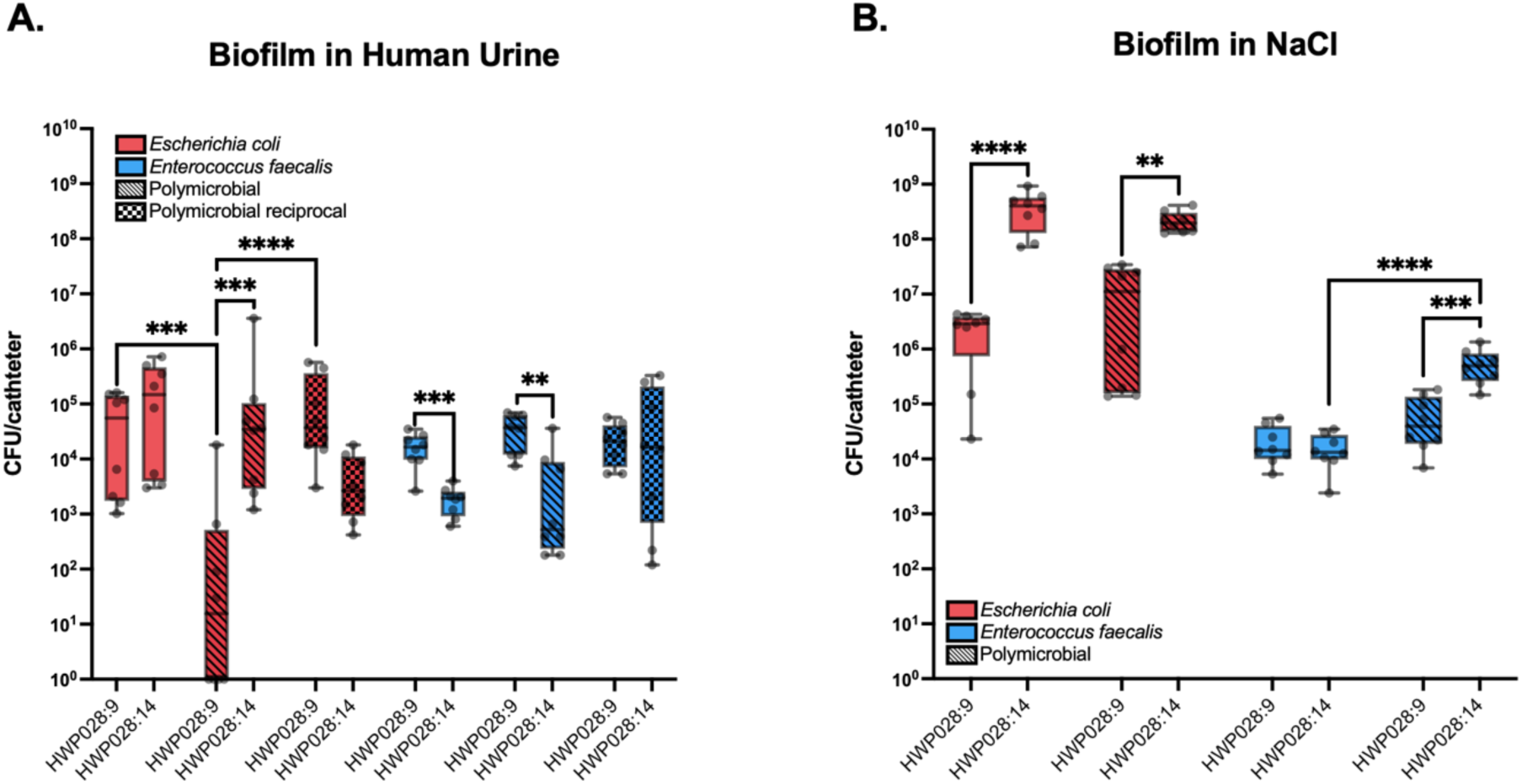
Biofilm formation in urine- and NaCl-supplemented medium. Biofilm formation on catheters was quantified as CFU per catheter after 48 h of growth for isolates obtained from two sampling timepoints (HWP028:9 and HWP028:14). *E. coli* is shown in red and *E. faecalis* in blue. Solid bars represent monoculture and lined bars represent polymicrobial conditions. Box-plot bars represent the min-max and median of eight biological replicates, and error bars indicate SD. Statistical comparisons were performed using an unpaired two-tailed t-test with Welch’s correction on log₁₀-transformed CFU values. **A.** Biofilm formed in BHI supplemented with 1:1 pooled human urine. Squared bars represent data where reciprocal pairing between HWP028:9 and HWP028:14 was performed. **B.** Biofilm formed in BHI supplemented with 1:1 0.9% NaCl.

To test whether this difference was due to *E. coli* or *E. faecalis*, we reciprocally paired each *E. coli* with the *E. faecalis* from the opposite time point. When *E. coli* HWP028:9 was paired with *E. faecalis* HWP028:14, the inhibition was lost, and *E. coli* recovered to monoculture levels (**Figure 5A**). *E. faecalis* HWP028:9 still reduced *E. coli* HWP028:14, but only slightly in comparison (approx. 19-fold), suggesting that *E. coli* HWP028:14 might also be less susceptible to inhibition. Reciprocally paired *E. faecalis* HWP028:14 reached densities comparable to HWP028:9, indicating that the stronger effect of HWP028:9 is a strain property rather than a result of better growth.

To test whether the strong inhibitory effects observed in urine reflected depleted nutrient content, we repeated the experiments in BHI diluted 1:1 with 0.9% NaCl. Instead, under polymicrobial conditions, both *E. coli* isolates maintained high biofilm counts with no significant reduction relative to monoculture (**Figure 5B**). *E. faecalis* monoculture biofilms were comparable between the two isolates, and under polymicrobial conditions *E. faecalis* HWP028:14 was significantly increased relative to its monoculture (**Figure 5B**). The loss of *E. coli* inhibition in this nutrient-reduced medium indicates that the urine effect is not explained by reduced nutrients alone.

### *E. faecalis* acidification matches the pH of clinical enterococcal bacteriuria

Clinical urine samples from ICU patients in our patient cohort with confirmed enterococcal bacteriuria displayed a lower pH (6.0) compared to bacteriuria-free ICU urine (6.5), urine from recovered post-ICU patients (6.5), and urine from ICU patients with non-enterococcal bacteriuria (7.0), as determined by clinical chemistry (multistix-7), suggesting that enterococcal colonisation is associated with urinary acidification *in vivo* (**Figure 6A**).

**Figure 6.**
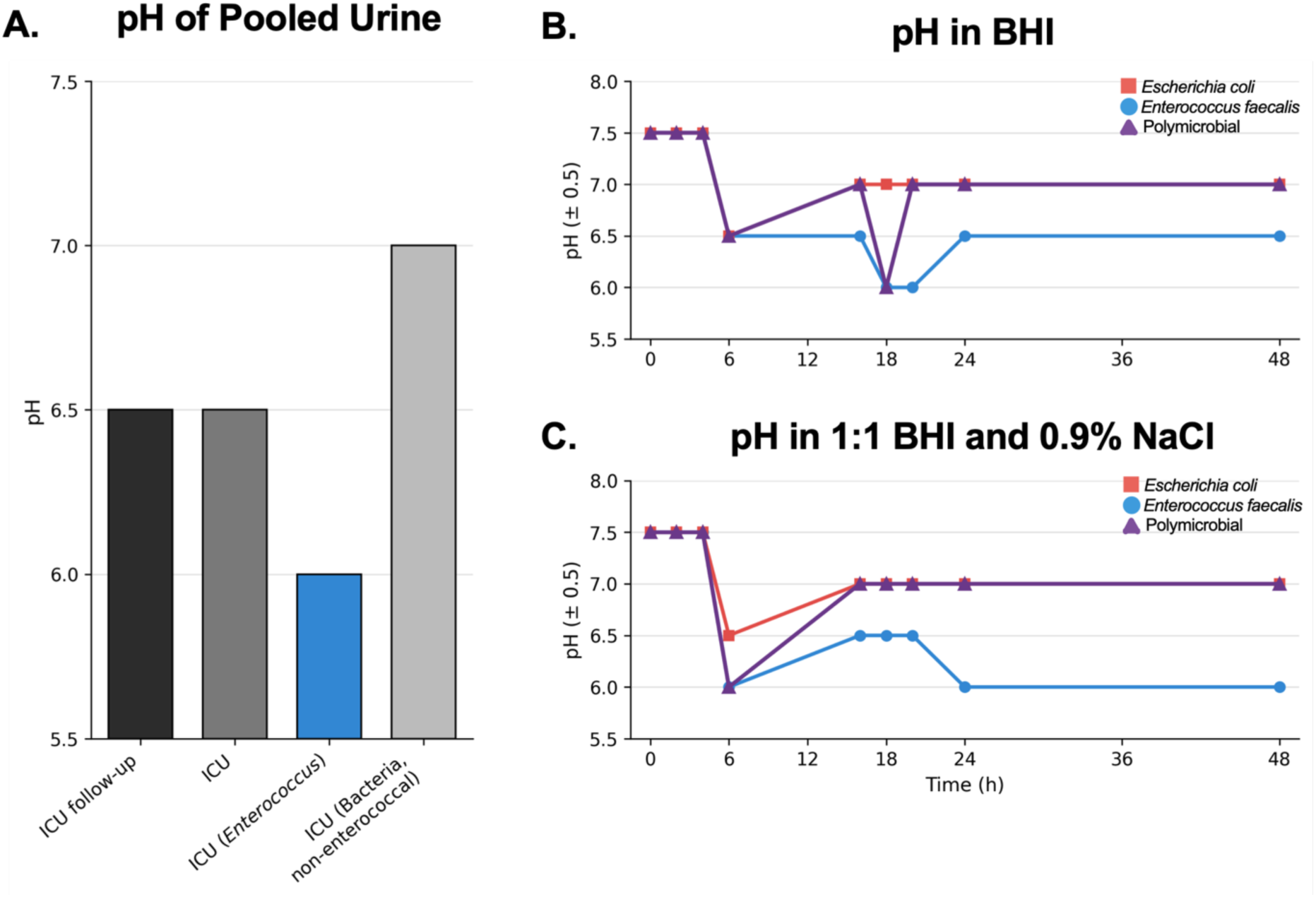
Clinical urine-pool pH and BHI culture pH dynamics. **A.** pH of pooled urine from four clinical categories: ICU follow-up (black), ICU without bacterial growth (dark grey), ICU with *Enterococcus* (blue), and ICU with non-enterococcal bacteria (light grey). Statistics unavailable. **B.** pH measured in BHI over 48 h for *E. faecalis* HWP028:9 monoculture (blue), *E. coli* HWP028:9 monoculture (red), and polymicrobial co-culture (purple). pH values are reported to the nearest 0.5 unit. Data points represent two identical biological replicates. **C.** pH measured in 1:1 BHI and 0.9% NaCl over 48 h. The same legend applies.

To relate biofilm outcomes to potential medium acidification, we monitored pH over 48 h in mono- and polymicrobial cultures of HWP028:9 in BHI and in BHI diluted 1:1 with 0.9% NaCl. The pH of the medium surrounding the polymicrobial biofilms (BHI, BHI-NaCl, and BHI-urine) was additionally assessed as snapshots at 0 h, 24 h (before media change), and 48 h (end of the experiment).

In BHI, uninoculated medium remained at pH 7.5 throughout. *E. coli* monoculture acidified to 6.5 by 6 h and recovered to 7.0 by 16 h, where it stayed. *E. faecalis* monoculture reached 6.5 by 6 h but declined further to 6.0 at 18 to 20 h before settling at 6.5. The polymicrobial culture held at 7.0 for most of the time course, apart from a transient dip to 6.5 at 6 h, and 6.0 at 18 h, coinciding with the *E. faecalis* nadir (**Figure 6B**). The BHI surrounding the biofilms followed the exact pattern of the time points, with pH 7.5, 7.0, and 7.0.

In BHI diluted 1:1 with 0.9% NaCl, *E. coli* monoculture again dipped to 6.5 at 6 h and recovered to 7.0, while *E. faecalis* monoculture fell to 6.0 by 6 h and fluctuated between 6.0 and 6.5. The polymicrobial culture dipped to 6.0 at 6 h but then held at 7.0 from 16 h onward, without the late dip seen in BHI (**Figure 6C**). The BHI-NaCl surrounding the biofilms again followed the pattern of the time points, with pH 7.5, 7.0 and 7.0.

In urine, the polymicrobial culture acidified strongly, falling from 7.5 to pH 5.0 by 24 h and remaining at 5.0 at 48 h even after the medium was replaced at 24 h, indicating sustained re-acidification.

### Catheter biofilms support strain-dependent survival and dispersal during TZP exposure

To assess whether the catheter-associated biofilm conferred tolerance to clinically relevant TZP concentrations, 48 h biofilms were exposed to 4 + 0.5 g/L TZP for 24 h and then allowed to recover for 48 h in antibiotic-free medium (**Figure 7A**). By broth microdilution in Mueller– Hinton, the TZP MICs for the strains were 0.5 mg/L for *E. coli* and 8 mg/L for *E. faecalis* (tazobactam fixed at 4 mg/L). MICs were higher in BHI, at 4 mg/L and 16 mg/L, respectively. Biofilms were exposed to the clinical 8:1 dosing ratio at approximately 1000-fold and 250-fold above the *E. coli* and *E. faecalis* BHI MICs.

**Figure 7.**
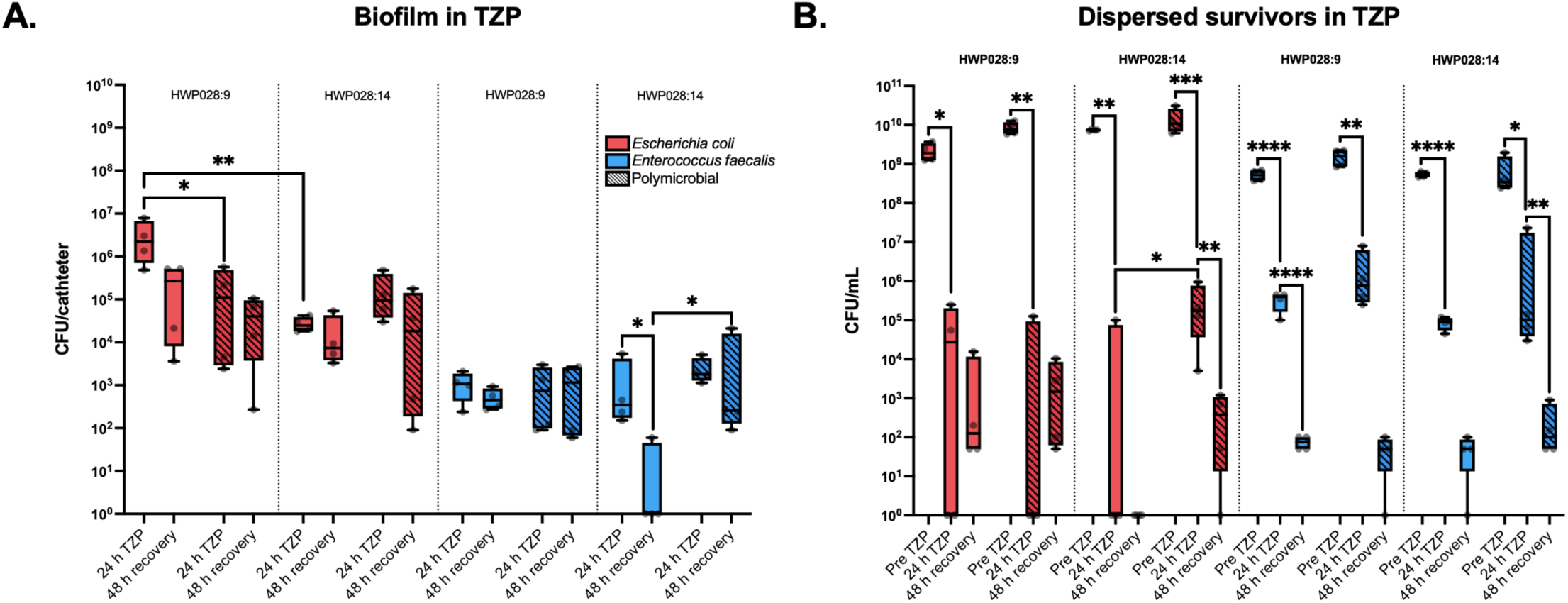
Biofilm survival and dispersal during TZP exposure and recovery. *E. coli* is shown in red and *E. faecalis* in blue. Solid bars represent monoculture and lined bars represent polymicrobial conditions. Box-plot bars represent the min-max and median of four biological replicates. Statistical comparisons were performed using an unpaired two-tailed t-test with Welch’s correction on log₁₀-transformed CFU values. **A.** CFU per catheter recovered from 48 h biofilms following 24 h TZP exposure and 48 h antibiotic-free recovery. **B.** CFU per mL recovered from the surrounding medium at each time point following media exchange, representing cells dispersed from the biofilm during each respective 24 h window. Pre TZP reflects basal dispersal from the biofilm at the 48 h timepoint in the absence of antibiotic.

Under TZP, *E. coli* HWP028:9 biofilms survived better than *E. coli* HWP028:14 in monoculture, reaching significantly higher CFU. Co-culture with *E. faecalis* reduced *E. coli* survival in a strain-dependent manner. For HWP028:9, polymicrobial *E. coli* was significantly lower than monoculture and remained so through recovery, whereas for HWP028:14 polymicrobial *E. coli* was comparable to, or slightly higher than, monoculture under TZP.

*E. faecalis* HWP028:9 remained low across all conditions, with no significant difference between monoculture and polymicrobial biofilm under TZP or at 48 h recovery (**Figure 7A**). For HWP028:14, monoculture *E. faecalis* fell significantly between 24 h TZP and 48 h recovery, whereas in co-culture it was maintained across the same interval and remained significantly higher than monoculture at recovery.

In parallel with the attached biofilm counts, the surrounding medium was sampled after each medium exchange, so that each sample represents only the cells released from the biofilm during that 24 h window (**Figure 7B**). Despite TZP concentrations far exceeding the MICs of both strains, dispersed *E. coli* were recovered at 24 h of TZP exposure in both mono- and polymicrobial conditions and in both isolate pairs, indicating that a subpopulation survived prolonged planktonic exposure to the antibiotic. The 48 h recovery sample reflects dispersal into fresh, antibiotic-free BHI. *E. coli* HWP028:14 monoculture released no culturable cells at recovery, indicating that this biofilm stopped shedding culturable cells after TZP exposure. In every other condition, dispersed *E. coli* remained detectable at recovery, though with considerable replicate variability, and generally at lower levels than at 24 h TZP, consistent with depletion or restriction of the dispersible reservoir. This reduction reached significance for HWP028:14 in co-culture but not for HWP028:9 in co-culture.

Dispersed *E. faecalis* were recovered at 24 h TZP across all conditions and both isolate pairs. For HWP028:9, both monoculture and polymicrobial dispersed CFU fell significantly between 24 h TZP and 48 h recovery and were comparable at recovery. For HWP028:14, monoculture dispersed CFU declined similarly, whereas the polymicrobial decline was less pronounced.

When non-biofilm-derived cells were exposed to this concentration planktonically in BHI broth, both species were killed, with no colonies recovered on agar plates even after 48 h. Dispersed biofilm survivors were additionally streaked onto BHI and MH agar plates, along with TZP E-tests, but no change in TZP MIC was observed. Survival under TZP therefore could not be attributed to intrinsic or acquired resistance.

### Dispersed TZP survivors show delayed regrowth consistent with a persister-like state

The dispersed survivors recovered under TZP raised the question of their physiological state and whether they could resume growth. Regrowth kinetics of dispersed HWP028:9 cells were measured against a standard curve of the unexposed control. Time to detection (TTD) scaled linearly with starting cell concentration for both species (*E. faecalis* R²: 0.97, *E. coli* R²: 0.99), each tenfold dilution delaying detection by approx. one hour (**Figure 8A, B**).

**Figure 8.**
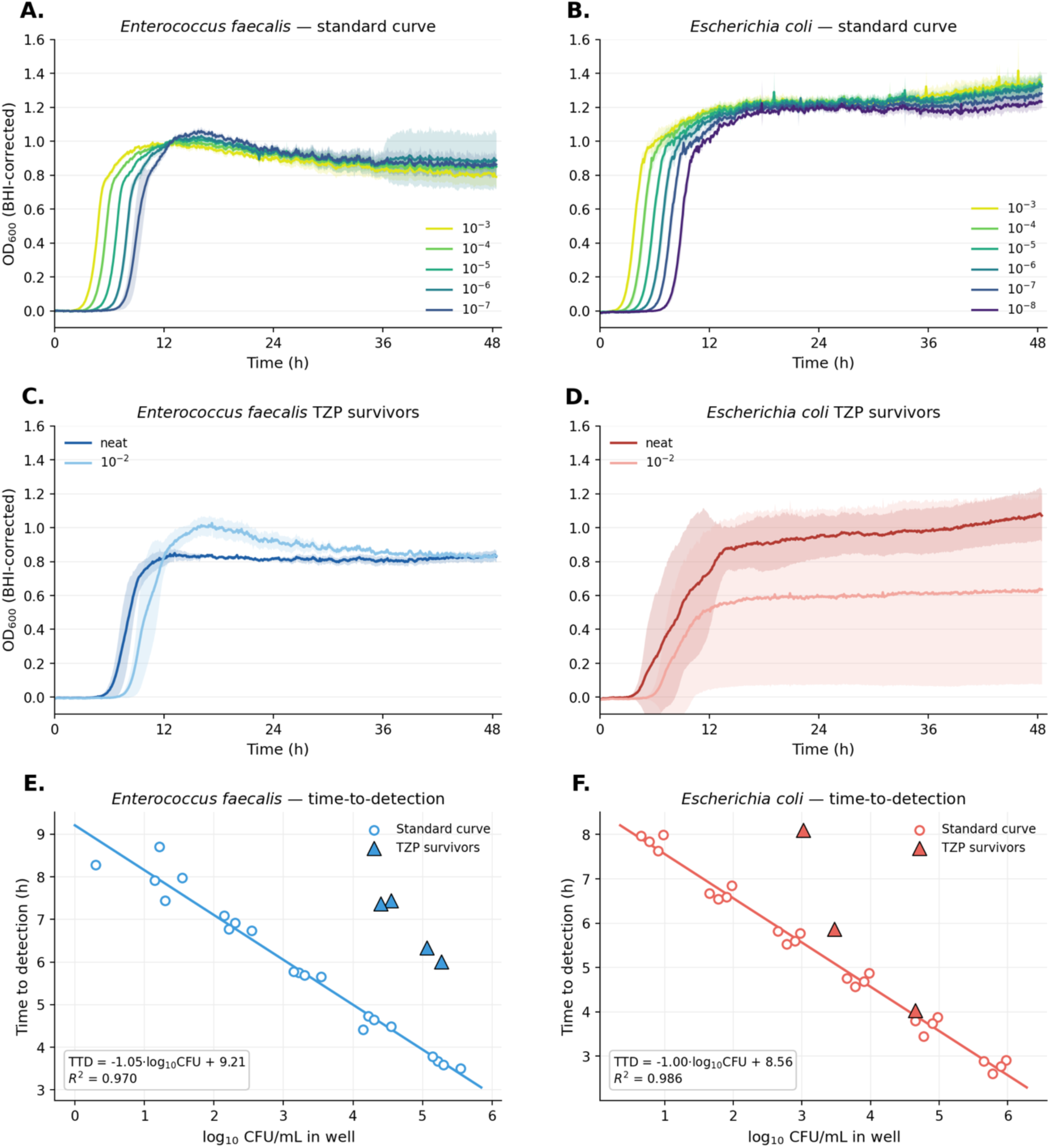
Regrowth kinetics of dispersed TZP survivors. Regrowth of dispersed HWP028:9 cells was monitored in a Bioscreen C. Optical density is shown after subtraction of the per-timepoint sterile-BHI background (OD₆₀₀, BHI-corrected). Growth curves are the mean of four biological replicates, with technical triplicates averaged within each replicate where present, and shaded regions denote one standard deviation. **A-B.** Standard curves of the unexposed control, serially diluted from 10⁻³ to 10⁻⁸ and colour-coded by dilution, for *E. faecalis* (A) and *E. coli* (B). The 10⁻⁸ dilution is omitted for *E. faecalis*, where only one replicate grew. **C-D.** Regrowth of TZP survivors loaded undiluted (neat) and at 10⁻², for *E. faecalis* (C) and *E. coli* (D). **E-F.** Time to detection (TTD), defined as the time to reach OD₆₀₀ of 0.1, plotted against the log₁₀ cell concentration in the well, for *E. faecalis* (E) and *E. coli* (F). Open circles represent standard-curve wells, the solid line represents the line of best fit by ordinary least-squares. Filled triangles represent TZP survivors plotted at their measured viable count. Survivors falling above the line reach detection later than unexposed cells of the same viable count. One *E. coli* replicate yielded no recoverable colonies and is not plotted.

TZP survivors regrew (**Figure 8C, D**) but reached detection later than unexposed cells of the same viable count (**Figure 8E, F**), following the signature of a persister subpopulation exiting dormancy before dividing. In *E. faecalis,* this delay was uniform, lagging by 2 to 3 h, equivalent to an apparent inoculum roughly 450-fold below the measured count. *E. coli* was more heterogeneous, with one biological replicate regrowing on the calibration line, one yielding no recoverable survivors, and the remaining having a lag phase of up to 2.5 h.

### Genomic analysis identifies an *esp* variant but no β-lactam resistance determinant

The pronounced phenotypic divergence between HWP028:9 and HWP028:14, despite recovery from the same patient and clonal complex, prompted three questions: (i) whether the paired isolates are genetically distinct, (ii) whether resistance genes underlie the TZP phenotypes, and (iii) whether either isolate encodes surface, persistence, or interaction features consistent with the polymicrobial biofilm behaviour.

Whole-genome sequencing confirmed that the paired *E. coli* isolates were indeed genetically indistinguishable at the nucleotide level, with no fixed single-nucleotide variants between HWP028:9 and HWP028:14. No plasmid replicons were detected in either isolate, and long-read sequencing revealed no additional contigs. The *E. coli* phenotypic divergence could therefore not be attributed to stable nucleotide differences or plasmid acquisition.

The paired *E. faecalis* isolates were highly similar but not identical. A repUS43 replicon marker was detected but mapped to the chromosome rather than a plasmid, and no plasmids were found in either HWP028:9 or HWP028:14. Comparative variant analysis identified four fixed nucleotide differences between the two time points. Two lay in rRNA regions, the 16S rRNA locus and the 23S–5S rRNA intergenic region. Because rRNA loci are multicopy and prone to mapping ambiguity, these two variants were treated cautiously and not taken as evidence of functional divergence.

The remaining two variants lay within *esp*, the gene encoding the enterococcal surface protein Esp. One mutation was synonymous, and one produced a Val1522Ile substitution. Among confidently interpretable coding differences, therefore, the only divergence between the *E. faecalis* timepoints fell within this biofilm-associated surface adhesin.

The *esp* gene lay within an IS6-family composite transposon that also carried the cytolysin operon and was flanked by *gls24* (stress response and persistence) and an araC-family transcriptional regulator (**Supplementary Figure 2**). Domain annotation predicted a C-terminal LPxTG-type cell wall anchor, comprising a non-canonical YPKTG motif followed by a hydrophobic segment and a positively charged tail, consistent with sortase-mediated anchoring. Both substitutions fell within the second antigen repeat domain, counted from the C-terminus. AlphaFold modelling suggested that Val1522Ile may be associated with a local conformational change in this region, with partial disruption of a β-sheet and emergence of a short α-helical segment (**Figure 9**). These predicted effects were local and should be read as hypothesis-generating rather than direct evidence of altered Esp function.

**Figure 9.**
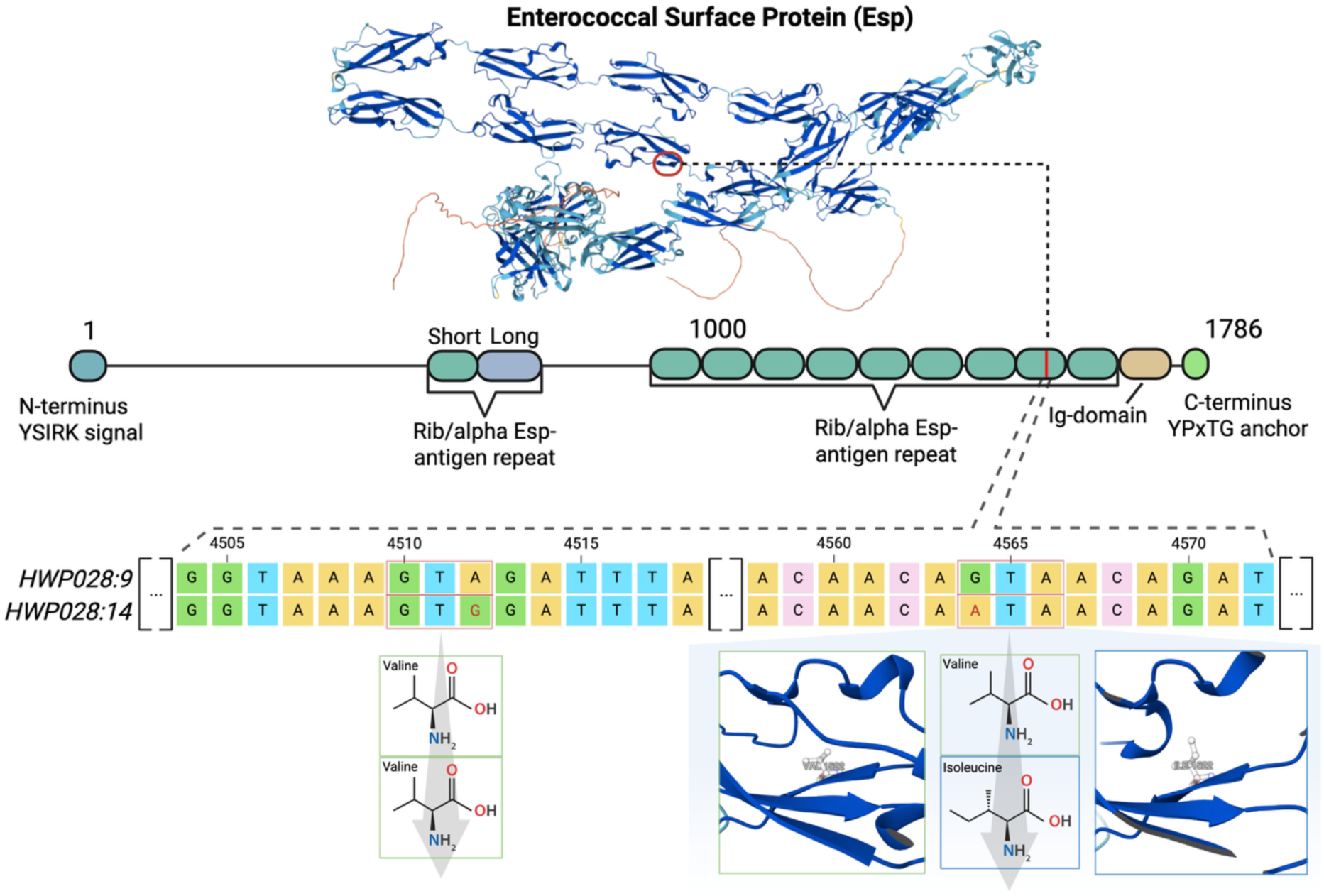
Comparative analysis of the Esp variant identified between day 9 and day 14 *E. faecalis* isolates. Comparative genome analysis identified two substitutions within the 5.37 kb gene encoding a predicted 1786 aa enterococcal surface protein (Esp). Domain annotation predicted a N-terminal YSIRK signal peptide, extensive internal Rib/alpha Esp antigen repeats, and a non-canonical YPKTG anchoring motif. Both detected mutations localized to the 2^nd^ antigen repeat (from the C-terminus) and comprised a synonymous substitution (Val1504Val) and a nonsynonymous substitution (Val1522Ile). Structural modelling using AlphaFold indicated that the Val1522Ile substitution is possibly associated with a localized conformational change within the repeat region, characterized by partial disruption of a β-sheet and emergence of a short α-helical segment, resulting in subtle local differences.

To test whether the TZP survival and recovery phenotypes could reflect antimicrobial resistance, resistance genes were predicted for the paired HWP028 isolates. The *E. coli* isolates carried no detectable resistance determinants (**Supplementary Table 1**). *E. faecalis* carried a limited resistome, with resistance predicted to trimethoprim (*dfrG*), tetracyclines (*tet(M)*), and macrolide–lincosamide–streptogramin antibiotics (*ermB* and *lsaA*) (**Supplementary Table 2**). No β-lactam resistance was predicted, and no known resistance-associated penicillin-binding protein (PBP) variant was identified. The same profile was present in both HWP028:9 and HWP028:14, indicating that the TZP survival phenotypes are unlikely to reflect acquired or intrinsic β-lactam resistance.

Consistent with the biofilm phenotypes, both species encoded extensive surface- and persistence-associated machinery (**Supplementary Tables 1 and 2**). *E. coli* carried a broad adhesin repertoire, including type 1 fimbriae and curli (*csgA*), consistent with attachment to and matrix formation on abiotic surfaces. *E. faecalis* encoded aggregation substance, the Esp and Ace adhesins, Ebp pili, sortase A, and pheromone-response systems, consistent with co-aggregation, surface persistence, and mixed-community biofilm formation. We identified the locations of both *E. faecalis* lactate production genes, *ldh1* and *ldh2*.

## DISCUSSION

This study set out to test two main hypotheses: (i) that *E. faecalis* aids the establishment of *E. coli* biofilms on urinary catheters, and (ii) that the resulting polymicrobial biofilm acts as the primary compartment protecting the community against clinically relevant β-lactam concentrations. Our results support the second hypothesis but reframe the first.

Contrary to the hypothesis, *E. coli* biofilms were inhibited in the presence of *E. faecalis*, and spent BHI from *E. faecalis* reproduced the inhibitory effect, indicating that the activity acts in solution rather than as a cell-cell-dependent residue. Spent medium from polymicrobial culture was less inhibitory than *E. faecalis* alone, which could reflect either consumption of the factor by *E. coli* or suppression of its production in the presence of *E. coli*. The fact that biofilm inhibition in BHI was observed only in the second round of incubation (48 h) also suggests that the inhibitory factor probably does not interfere with biofilm initiation or has not yet been produced at an early stage. The identity of the factor remains open, but candidates supported by the genome include organic acids, as common *E. faecalis* bacteriocins, including the identified cytolysin, have been shown not to act on *E. coli* [59,60].

More likely, *E. faecalis* shapes the local environment by e.g. local acidification, primarily through lactate dehydrogenase [61]. *E. coli* is known to be inhibited and permeabilized by lowered pH, and particularly so by lactic acid [62]. Even when not lethal, *E. coli* must spend energy to neutralize local acid [63,64]. Even L-lactate on its own might have an inhibitory effect [65].

For these reasons, and because acidification has previously been proposed as an inhibitory mechanism against other Gram-negatives, we set out to investigate how pH might relate to our biofilms. In well-buffered BHI, *E. faecalis* lowered the pH sharply (to pH 6.0) but only transiently, first at 18 h, whereas in 1:1 BHI-NaCl, the sharp dip occurred early (6 h) and later stabilized at the pH of the *E. coli* monoculture.

One hypothesis is that pure BHI, with its greater buffering capacity and fermentable load, delayed net acidification by prolonging acid production into the later growth phase, before *E. coli* acid-resistance mechanisms were activated. In contrast, acidic saline dilutes and partially consumes the phosphate buffer and reduces fermentable substrate, potentially causing an earlier pH drop without a later acidogenic phase.

The BHI-NaCl medium was the only medium in which *E. coli* biofilm was not inhibited by *E. faecalis*. While it is possible that the later 18 h dip observed in BHI interferes with or stunts *E. coli* biofilm maturation, such a stress response should arguably be released during the BHI media exchange at 24 h. While purely speculative, it might be that an early acid pulse shapes the culture towards biofilm production, while a late acid pulse shapes a pre-established biofilm towards dormancy, or it interrupts biofilm maturation at a crucial stage. The pH measured in our media is sufficient to initiate mild to moderate acid resistance (AR) systems in *E. coli*, which are known to induce growth arrest and elongation [66,67].

In poorly buffered BHI-urine, the polymicrobial culture fell to pH 5.0 by the 24th hour, consistent with the lower pH measured in clinical urine from ICU patients with enterococcal bacteriuria. Interestingly, at this lowered pH, *E. coli* must activate AR3 and AR5 [68], leading to secretion of agmatine and putrescine, both of which *E. faecalis* efficiently metabolizes [69]. Acidification therefore remains a potentially inhibitory factor, and future studies might investigate whether *E. faecalis* can directly benefit from stimulating an acid response, as previously suggested [70].

In urine, the two isolate pairs, recovered before and after antibiotic exposure, behaved differently, and reciprocal pairing localized this difference to the *E. faecalis* partner. The early *E. faecalis* (HWP028:9) was strongly inhibitory, whereas the later *E. faecalis* (HWP028:14) was not, despite reaching comparable densities. A smaller, secondary contribution came from the *E. coli,* where the HWP028:14 isolate was somewhat less susceptible, consistent with being a better biofilm former in BHI. Genomically, the *E. coli* isolates were genetically indistinguishable. Hence, their phenotypic divergence must have a non-genetic basis. In contrast, the *E. faecalis* strains differed at only one coding position, a Val1522Ile substitution in the second antigen repeat of the surface adhesin Esp.

Esp is a large, cell-wall-anchored surface protein attached to the peptidoglycan and projecting from the cell as a periscope built from tandem antigen repeats [71,72]. It has been repeatedly associated with adhesion, biofilm formation, and urinary tract colonization, contributing to bladder persistence. The substitution itself, however, is conservative. Valine and isoleucine are both small, branched, hydrophobic residues, and any potential relevance should therefore lie in structural context. Within a tightly packed repeat fold, even a near-isosteric change can affect local packing, consistent with the conformational change predicted by our structural modelling. Esp assembles into amyloid-like fibers under acidic conditions that stabilize the biofilm matrix [73], and by sealing internal microenvironments, Esp may hypothetically modulate outward diffusion of metabolic byproducts. Although our substitution lies within the repeat region, it could contribute to higher-order assembly.

The substitution also lies within a composite transposon, alongside *gls24*, a stress regulator whose disruption drives shifts in enterococcal chain length and virulence, and an AraC-family regulator governing numerous genes involved in pH homeostasis and biofilm formation [74,75]. The substitutions may therefore mark a broader, regulatory difference, attuning the HWP028:14 isolate toward a phenotype that permits *E. coli* to mature within the shared biofilm. These interpretations remain speculative, but given that *E. faecalis* biofilm yielded substantially higher CFU in the presence of *E. coli*, adaptation might ultimately be beneficial for them both.

How the two species co-localized in the bladder and on the catheter at the same time, despite apparent initial antagonism, can partly be explained by co-aggregation. We showed that *E. coli* and *E. faecalis* co-aggregated in suspension before even attaching to the catheter surface, with *E. coli* rods physically embedded, alive, within sedimenting *E. faecalis* aggregates. Several features of *E. faecalis* could actively drive this association.

As a producer of autoinducer-2 (AI-2), a universal LuxS-dependent quorum-sensing signal, *E. faecalis* has previously been shown to recruit *E. coli* via chemotaxis, thereby increasing the probability of collision [76]. Our *E. faecalis* additionally carried aggregation substance (AS), a pheromone-inducible surface adhesin that mediates cell-cell clumping. AS drives *E. faecalis* into large self-aggregates and greatly increases the target size and local concentration of AI-2 [39,77,78]. The resulting clumps might act as sedimenting nets that entrap passing *E. coli*, raising the probability of capture far above that expected for freely dispersed cells. AS additionally increases cell-surface hydrophobicity, promoting nonspecific hydrophobic adhesion through which *E. coli* would become bound to, and embedded within, the aggregate. Because catheter colonization proceeds largely along the extraluminal catheter–urethra interface, ascending co-aggregates could be co-introduced [31].

On the catheter itself, the two species formed four distinct microenvironments in which *E. faecalis* abandoned its discrete cluster morphology for a dispersed, EPS-producing form associated with *E. coli*, something which we have not seen reported elsewhere. Why *E. faecalis* would adopt this configuration remains unexplored. A dispersed, EPS-embedded form might maximize the interface over which the two species can perform nutrient exchange, as *E. faecalis* is a documented metabolic donor of L-ornithine (additionally acting as a substrate for AR5 putrescine secretion), which *E. coli* uses to upregulate enterobactin biosynthesis for iron acquisition [61,79]. Switching from a discrete cluster might also represent a more aggressive strategy, focusing on suppression of *E. coli* by expanding across the catheter surface before *E. coli* otherwise uses it to establish a single-species biofilm. The dependence may also run the other way. *E. coli* is an efficient colonizer equipped to attach even to medical-grade silicone, while previous studies have argued that *E. faecalis* might need a fibrinogen coating for proper biofilm development [37]. An established *E. coli* biofilm might hence present a far more favourable biological substrate for such a biofilm to mature. Read together, *E. faecalis* neither excludes nor merely tolerates *E. coli*, but rather appears to exploit it.

The survival of both species under TZP was a property of the biofilm and not of resistance. Neither isolate carried any β-lactam resistance genes or a resistance-associated PBP variant, and planktonic cells exposed to the same concentration in broth were killed, with no survivors even after 48 hours of recovery. Biofilm-derived cells in suspension survived exposure hundreds-fold above the MIC, but when survivors were tested against TZP using E-tests, no changes compared to their planktonic counterpart were observed. In monoculture, *E. coli* HWP028:14 biofilm was more susceptible to TZP than the earlier isolate, the opposite of what would be expected if it had acquired tolerance during its 11-day survival in the patient. Its survival is therefore better explained by residence in the polymicrobial community than by any intrinsic change. Consistent with this, protection became reciprocal in the latter pair, as *E. faecalis* HWP028:14 monoculture collapsed after antibiotic removal, whereas in co-culture with *E. coli* it was maintained. While it is compelling to suggest that the Esp substitution might have provided a trade-off between monoculture resilience and polymicrobial adaptation, future studies will have to confirm the exact role of Esp in these settings.

The cells released from the biofilm in the presence of TZP were physiologically distinct. Measuring growth kinetics following antibiotic exposure, dispersed survivors reached detection later than unexposed cells with the same viable count, the signature of a persister subpopulation that must exit dormancy before resuming growth [80,81]. In *E. faecalis,* this lag was uniform across replicates, corresponding to an apparent inoculum several hundred-fold below the measured count. That the *E. coli* HWP028:14 monoculture retained viable attached cells within its biofilm, yet yielded no culturable cells in the surrounding medium at recovery, indicates that survival was confined to the biofilm compartment.

Whether these survivors reflect persistence that is specifically biofilm-dependent remains an unresolved question, since persister formation is classically observed in planktonic, stationary-phase populations as well as in biofilms. A mechanism that could reconcile this with the broader literature is the phenotypic memory of biofilm-derived cells, whereby cells might retain a biofilm-associated tolerant state after dispersal [82–84]. This would help explain that cells recovered from the surrounding medium, already detached from the biofilm, still displayed persister kinetics whereas planktonic cells did not.

Together, these findings offer a mechanistic account of the epidemiological pattern that motivated the study, in which broad-spectrum β-lactam therapy favours enterococci while suppressing susceptible uropathogens. Urinary infections under such therapy are often attributed to resistance, but we have shown here that resistance is not required. Our isolates were fully susceptible at the concentrations used, and their survival instead reflects biofilm-dependent persistence. *E. coli* is commonly indicated as a driver and protagonist, but we argue that *E. coli* might in fact be recruited into shared biofilms, orchestrated by *E. faecalis* aggregation. As a future direction, it would be useful to investigate whether such biofilms could form on the catheter during ascension of the catheter–urethral interface, allowing cells to reach the catheterized bladder in a dormant, potentially co-aggregated state. In such circumstances, ongoing antibiotic therapy would not be sufficient to prevent colonization, which would only become apparent after treatment is discontinued.

There are several limitations to address. The study centres on a single patient’s isolate pairs, and although this longitudinal, before-and-after design is a strength, the generality of the interactions will require follow-up experiments with additional strain pairs. Bulk colony counts averaged across the four identified biofilm microenvironments revealed by SEM cannot resolve which compartment survives a stressor, such as TZP, and which is eliminated. Absolute biofilm and dispersed counts are affected by the fact that our assay does not identify viable-but-non-culturable state cells. Finally, biofilm formation was unexpectedly enhanced in nutrient-diluted, salt-supplemented medium relative to full BHI, an effect outside the present scope that merits separate investigation.

In conclusion, a clinically susceptible *E. faecalis*–*E. coli* community sustained on a urinary catheter survives supra-MIC β-lactam therapy not through resistance but through biofilm-dependent persistence and interspecies adaptation. *E. faecalis*, despite suppressing *E. coli* independent biofilms, captures it into this community and, under antibiotic stress, comes to depend on it in return. Recognizing these communities as reservoirs of tolerance rather than resistance reframes how persistent and recurrent catheter-associated infections might be prevented and treated.

## ETHICAL STATEMENT

The samples used in this study are part of the PronMed study, approved by the Swedish National Ethical Review Agency (Dnr 2017/043, with amendments 2020-01623, 2020-02719, 2020-05730, and 2021-01469; and Dnr 2022-00526-01) and registered at ClinicalTrials.gov (NCT03720860). The study was conducted in accordance with the Declaration of Helsinki and its subsequent revisions. Informed consent was obtained from each patient or their next of kin.

## ACKNOWLEDGEMENTS

The authors thank Karin Hjort and Greta Zaborskyte for their input in establishing and optimizing the early catheter biofilm system, Nikolaos Kavalopoulos for suggesting the sealing of catheter inflation lumina and for contributing silicone reagents, Anders Larsson for assistance with the clinical-chemistry analysis of patient urine samples, Xiguo He for help in validating the assessment of urinary piperacillin concentrations, and Linglu Hong and the BioVis platform at Uppsala University for advice on SEM preparation and for contributing fixation reagents. We thank Michael Hultström, Robert Frithiof, and Miklos Lipcsey for the organization, establishment, and overall responsibility of the PronMed study, and Erik A. Danielsson, Joanna Wessbergh, and Elin Söderman for their contributions to patient recruitment and sample collection. We are also grateful to students Penina Thell, Linnea Persson, and Taoran Zhang for their contributions during the project. This study was funded by WISE-WASP (**WX**: LiU-2023-00139, Knut and Alice Wallenberg Foundation) and the Swedish Research Council (**Francesco D’Elia**: 2025-04795, **HW**: 2024-03665). The funder played no role in study design, data collection, analysis and interpretation of data, or the writing of this manuscript.

## AUTHOR CONTRIBUTIONS

**PK**, **HW**, and **JJ**: conceptualization. **PK**: writing original draft. **PK**, **MS**, **CB**, **HZ**, **HA**: data curation. **PK**, **EJ**: formal analysis. **HW**, **WX** and **JJ**: funding acquisition. **PK**, **HW**, and **JJ**: investigation. **PK**, **CB**, **HW** and **JJ**: methodology. **PK**, **HW**, **WX** and **JJ**: project administration. **HW**, **WX** and **JJ**: validation. All authors contributed to the interpretation of results and critical review of the manuscript and had access to the data, except for identifiable clinical data to which access was restricted to those acquiring and analyzing it.

