## Supplementary for "Polymicrobial catheter biofilms sustain susceptible *Enterococcus faecalis* and *Escherichia coli* during β-lactam treatment"

### SUPPLEMENTARY MATERIAL

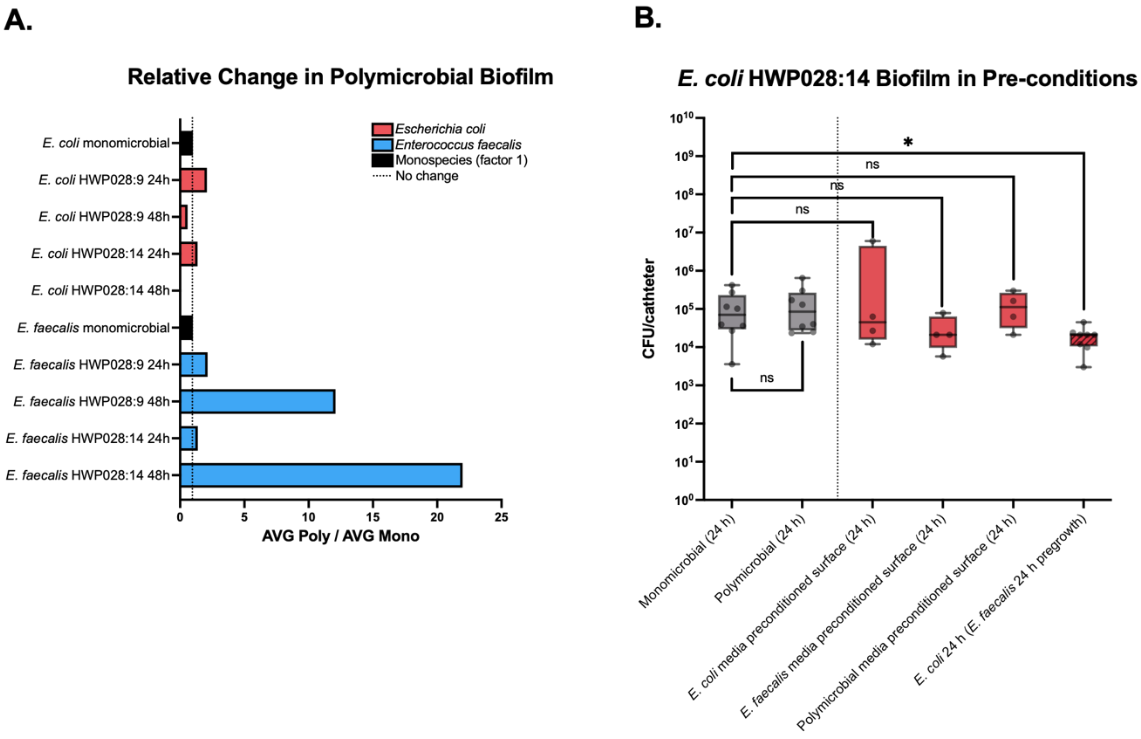


**S. Figure 1. *E. faecalis* enhances its own biofilm in polymicrobial culture. A.** Relative change in biofilm formation in polymicrobial versus monospecies conditions, expressed as the ratio of mean polymicrobial to mean monospecies biofilm density (CFU per catheter). Each species' monoculture value is set to 1 (black), and ratios above 1 indicate greater biofilm in polymicrobial culture. Bars are ratios of group means and are descriptive only, hence no statistical testing was applied. **B.** Catheter segments were incubated in cell-free spent medium from *E. coli*, *E. faecalis*, or polymicrobial cultures for 24 h to allow surface adsorption of soluble constituents. The spent medium was then removed, and the preconditioned segments were submerged in *E. coli* inoculum for a further 24 h before biofilm quantification. Data from Figure 1AB is reused for comparative reasons (grey).

**S. Table 1. Molecular characterization of *Escherichia coli* HWP028:9**

| Category | Feature | Result | Biological relevance |
|---|---|---|---|
| **Core lineage typing** | cgMLST | cgST 116433 | - |
| | CH type | fumC13–fimH41 | - |
| **Serotype** | O antigen | O4 | LPS surface structure, host interaction, surface properties |
| | H antigen | H1 | Flagellar motility |
| **Primary adhesion systems** | Type 1 fimbriae | fimH41 | Early surface attachment, biofilm initiation |
| | P fimbriae | *papA, papC* | Uroepithelial adhesion, biofilm initiation |
| | S/F1C fimbriae | *sfaD, focC* | Host tissue binding, biofilm initiation |
| | Additional adhesins | *iha, fdeC, yeh/yfc* cluster | Surface colonization |
| | Invasion determinant | *tia* | Host cell interaction |
| **Biofilm matrix factors** | Curli production | *csgA* | Extracellular matrix formation |
| **Iron acquisition systems** | Aerobactin | *iutA, iuc* | Growth in iron limitation |
| | Yersiniabactin | *fyuA, irp2* | Growth in iron limitation |
| | Heme uptake | *chuA* | Iron scavenging |
| | Salmochelin receptor | *iroN* | Nutrient acquisition |
| | Additional transport | *sitABCD, ireA* | Stress fitness |
| **Toxins/host interaction** | α-hemolysin | *hlyA* | Host cell damage |
| | Uropathogenic protein | *usp* | UTI-associated factor |
| | Immune modulation | *tcpC* | Host immune interference |
| | Genotoxin island (*pks*) | *clbB* (colibactin) | Host interaction |
| | Outer membrane protease | *ompT* | Defence/virulence |
| **Competition factors** | Microcin system | *mch* genes | Interbacterial competition |
| | Colicin | *cea* | Competitive fitness |
| **Stress response** | Capsule genes | *kpsE, kpsMII* | Immune evasion, biofilm protection |
| | Serum survival | *iss* | Host persistence |
| | Stress resistance | *terC* | Environmental tolerance |

**S. Table 2. Molecular characterization of *Enterococcus faecalis* HWP028:9**

| Category | Feature | Result | Biological relevance |
|---|---|---|---|
| **Serotype** | - | ST16 | - |
| **Antimicrobial resistance** | Trimethoprim resistance | *dfrG* | Antifolate resistance |
| | Macrolide–lincosamide resistance | *erm(B)* | MLS resistance phenotype |
| | Tetracycline resistance | *tet(M)* | Ribosomal protection |
| | Streptogramin/lincosamide resistance | *lsa(A)* | Intrinsic resistance mechanism |
| **Primary adhesion systems** | Aggregation substance | agg | Cell–cell aggregation, conjugation, biofilm formation |
| | Surface adhesin | *esp* | Biofilm formation, persistence |
| | Collagen adhesin | *ace* | Host tissue binding |
| | Endocarditis antigen | *efaA/fimA* | Host colonization |
| | Sortase | *srtA* | Surface protein anchoring |
| | Endocarditis biofilm pili | *ebpA, ebpC* | Biofilm formation on surfaces |
| **Toxins/host interaction** | Cytolysin operon | *cylA, cylB, cylM, cylI, cylL, cylR* | Host cell damage, interbacterial competition |
| | Hyaluronidase | *hylA* | Tissue invasion potential |
| | Hemolysin III | *yqfA* | Host cell damage |
| | Virulence regulator | *elrA* | Pathogenicity control |
| **Pheromone response** | Sex pheromone systems | cAM373 (camE), cCF10, cOB1 | - |
| **Stress response** | Oxidative stress defence | *tpx* | Survival under stress |
| | Metal resistance | *cad* | Environmental persistence |

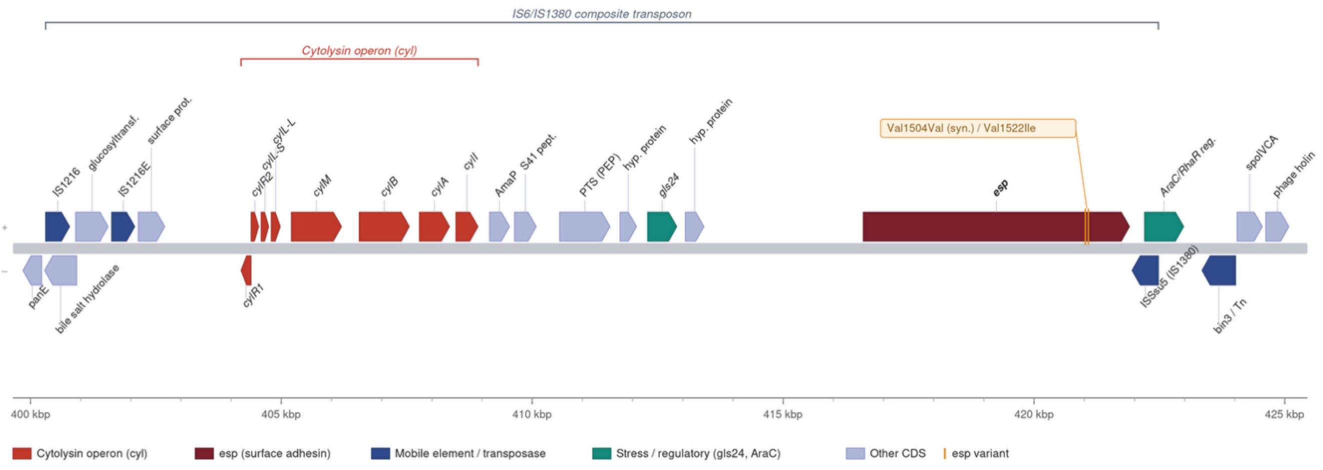


**S. Figure 2. *esp* localizes inside an IS6-family composite transposon.** Part of *E. faecalis* HWP028:14 chromosome illustrating the IS6 transposon (blue) containing the *cyl*-operon (light red), regulatory response elements (green) and the *esp* gene (dark red), with two identified substitutions (orange).

#### Piperacillin–Tazobactam Concentrations for Biofilm Experiments

All isolates in this study originated from an adult ICU patient with severe COVID-19 at Uppsala University Hospital, who uniformly received intravenous TZP at a total dose of 12 g piperacillin and 1.5 g tazobactam per 24 hours.

Piperacillin is primarily eliminated unchanged by the kidneys (60–70%), with about 10–15% metabolized by the liver to inactive desethylpiperacillin (which is also excreted in urine) and 5–10% excreted in bile. Spontaneous hydrolysis of piperacillin in urine is minimal, at 5% or less, under normal clinical conditions. Tazobactam is eliminated mainly by renal excretion (about 80% unchanged), with 20% metabolized (to inactive form M1) by the liver and minimal biliary or spontaneous urinary hydrolysis (Pfizer, LAB-0446-7.0).

According to Wallenburg et al. (2022), critically ill patients receiving 12 g/24 h piperacillin reach median 24-hour urine piperacillin concentrations between 3000 mg/L and 6000 mg/L (creatinine clearance 70–116 mL/min). These findings are corroborated by our cohort's renal function, as detailed in Larsson et al. (2022), where median estimated GFR (by cystatin C and creatinine-based equations) was 70–83 mL/min. Applying the Wallenburg PK model, these values yield similar piperacillin urine concentrations.

$$C_{urine} = \frac{Dose\ per\ 24h\ \times\ Fraction\ excreted\ unchanged\ at\ 24h \times\ (1 - Fraction\ hydrolysed)}{Urine\ volume\ per\ 24h}$$

While clinical breakpoints and dosing recommendations for TZP are determined based on plasma pharmacokinetics and pharmacodynamic targets (e.g., achieving 100% fT>MIC in plasma for pathogens with MIC ≤8 mg/L), urine concentrations in patients receiving standard dosing become several thousand mg/L, far exceeding these breakpoints. As a result, organisms classified as "resistant" in plasma-based testing may still be rapidly eradicated from the urinary tract, provided the infection remains localized to the urine and does not involve tissue or bloodstream invasion.

For piperacillin, with a daily dose of 12 g, we assume that 70% is excreted unchanged, 5% hydrolysed, and a urine output of 1.5–2.0 L, yielding urine concentrations of piperacillin between approx. 4,000 and 6,000 mg/L.

### Catheter-based polymicrobial biofilm assay on medical-grade silicone urinary catheters

This protocol is written to be self-contained. Volumes, concentrations, incubation times, and instrument settings are given explicitly. The 3D-printing files and the original description of the FlexiPeg platform are required accessory materials and are cited where used.

#### 1. Principle and overview

Segments of a clinical silicone Foley catheter are mounted vertically in a 3D-printed modular tray so that each segment is held in one well of a 96-well microtiter plate. Segments are submerged in a defined bacterial inoculum and incubated statically, allowing biofilm to develop on the true luminal and abluminal catheter surfaces. The modular tray allows the whole set of segments to be transferred between plates, so mature biofilms can be moved through medium changes, alternative media, or antibiotic exposure without disturbing the attached community. After a defined incubation, biofilm is quantified from the catheter surface, the surrounding medium is sampled in parallel, or segments are processed for scanning electron microscopy (SEM).

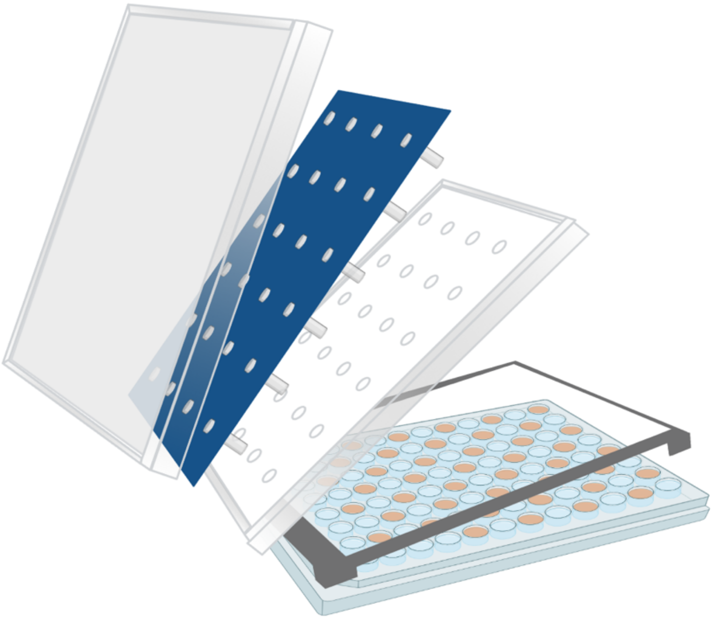

The model is designed to reproduce catheter material, retained luminal geometry and restricted gas exchange. Catheter segments are reusable after a defined cleaning cycle.

##### Assay timeline overview

**One-time:** prepare and seal catheter segments.

- **Day −1:** streak strains and prepare fresh media.
- **Day 0:** prepare inoculum, mount sterile catheter module, submerge segments, incubate 24 h.

- **Day 1:** transfer module to fresh medium/condition on a 2 mm insert (or read out) and incubate a further 24 h. A mature biofilm in this protocol is 2 × 24 h.
- **Day 2 onward:** read out (catheter CFU, broth CFU, crystal violet, or SEM), or continue into a variant module (conditioned medium, sequential colonization or antibiotic exposure and recovery).

### 2. Materials and equipment

#### 2.1 Catheters and the modular system

| Item | Specification |
|---|---|
| Silicone Foley catheter | Teleflex Brilliant Plus Aquaflate, 100% silicone, 12 Ch, 40 cm, 10 mL balloon, cylindrical tip. |
| FlexiPeg modular tray set | 3D-printed from the STL files accompanying Zaborskyte et al. 2021: silicone-mat mould (used to cast the silicone-mat), 96-well lid, peg-lid top, and the 2 mm elevation insert. |
| Silicone mat | Cast from the mould STL using the same elastomer as below. Holds catheter segments upright over a 96-well plate. |
| SYLGARD 184 elastomer kit | Base:curing agent 10:1. Used to seal catheter inflation lumina. Cures 4 h at room temperature or 1 h at 60 °C. Store uncured mix at −20 °C to delay curing. |
| 96-well flat-bottom plates | Sterile, standard. One plate holds up to 6 conditions × 4 biological replicates plus controls. |
| 27 G cannula + syringe | For injecting elastomer into the inflation lumen. |

#### 2.2 Media, reagents, consumables

| Item | Specification |
|---|---|
| Brain Heart Infusion (BHI) | Broth and agar (Oxoid). Prepared fresh weekly, autoclaved in UV-opaque (foil-wrapped) glass, cooled and stored at 4 °C in the dark, used within 5 days. |
| 0.9% NaCl (saline) | For inoculum standardisation. |
| Phosphate-buffered saline (PBS) | Sterile, for washing and serial dilution. |
| Brilliance UTI Clarity agar | Oxoid/Thermo Fisher. Chromogenic differentiation of *E. coli* and *E. faecalis* in polymicrobial platings. |
| Crystal violet | 1% aqueous stock, diluted 1:20 to 0.05% working solution. 95% ethanol for solubilisation. |
| SEM fixatives / buffers | Sörensen's phosphate buffer, 2.5% glutaraldehyde + 1% paraformaldehyde, graded ethanol 30–99.5%. |
| Piperacillin and tazobactam | Sterile-filtered aqueous stocks (40 g/L piperacillin, 5 g/L tazobactam) for the antibiotic module. |
| Contrad 70 | Fisher. 5% cleaning solution for catheter reuse. |
| Isopropanol, ethanol | Cleaning. Tweezer sterilisation and insert storage. |
| Atmosphere-altering sachets | CampyGen Compact (Thermo Fisher, CN0020C) for microaerophilic incubation. |

##### 2.3 Equipment

- Nephelometer with 0.5 McFarland standard.
- Autoclave, 37°C, 60°C and 50°C incubators.
- Airtight incubation boxes and 1 L gas-barrier bags (for microaerophilic incubation).
- Multi-tube vortex (biofilm resuspension and crystal-violet solubilisation).
- Microplate reader (e.g., Thermo Scientific Multiskan FC). OD 620 nm for broth, OD 540 nm for crystal violet.
- Sputter coater (e.g., Polaron SC7640) and field-emission SEM (e.g., Zeiss Merlin), SEM pin stubs, mount holder, conductive carbon tape (SEM performed with trained facility staff).

### 3. One-time catheter preparation

##### 3.1 Sealing the inflation lumen

Sealing the balloon-inflation lumen removes a small blind channel that otherwise traps liquid and introduces washing/staining variability. It also better reflects the in-use catheter. In the standard set-up, all inflation lumina are sealed.

1. Cut off both ends of the full catheter to expose the inflation tube at each end.
2. Prepare SYLGARD 184 by mixing base and curing agent 10:1.
3. Using a syringe fitted with a 27 G cannula, inject the elastomer into the inflation lumen until the entire channel is filled. Apply steady, moderate pressure to avoid trapping air bubbles.
4. Cure at 60 °C for 60 min (or 4 h at room temperature).

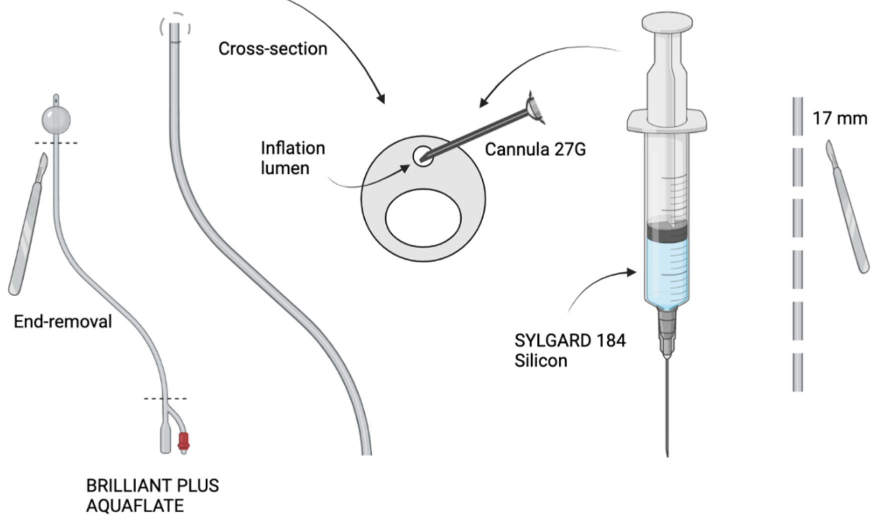

**Note.** Cured silicone cannot be removed from surfaces or glassware. Dedicate consumables to this step and wear gloves.

##### 3.2 Cutting segments

1. Cut the catheter into 17 mm segments.
2. Put each new segment through one full cleaning cycle before first use, to remove cutting debris and standardise handling.

**Note.** Segment length sets the liquid volume the lumen can hold, and therefore the assay volumes. A 17 mm segment holds roughly 60–70 µL. Segments must be ≥17 mm to fit the plate wells. Keep lengths constant across an experiment.

#### 4. Module assembly and sterilisation

Perform at least one day before the experiment.

1. Place the silicone mat on top of the FlexiPeg tray.
2. Rest the mat-and-tray module over a flat-bottom 96-well plate (the plate is only an alignment aid while pushing the segments down).
3. Insert catheter segments into the mat, all in the same orientation. The lower end of the catheter should touch the bottom of the well.
    a. Keep every segment in the same rotational orientation (e.g., the cross-sectional white indicator marking facing up, the thicker sealed-lumen side facing the same way) so that rubbing marks from the mat and tweezers always fall on the same region and do not confound the biofilm-exposed surface.
4. Inspect from underneath and straighten any slanted or bent segments.
5. Place the module on a protective support (e.g., an inverted pipette-tip-box lid) so the segments are shielded, then seal inside an autoclave bag.
6. Autoclave at ≥120°C for 15 min.
7. Dry the bagged module at 50 °C (3–4 h or overnight) until completely dry.

**Critical.** When later removing the module from the bag, work in a decontaminated, uncluttered space with fresh gloves that have touched nothing else. Contamination introduced here compromises the whole plate.

### 5. Bacterial culture and inoculation

#### 5.1 Preparing and standardising the inoculum

1. The day before the experiment, streak each strain onto any non-selective/non-indicative agar and incubate. Always use freshly grown colonies.
2. Calibrate the nephelometer against the 0.5 McFarland standard.
3. Suspend fresh colonies in 5 mL 0.9% NaCl and adjust with saline or additional colonies to 0.5 McFarland (≈ $10^8$ CFU/mL). Screw-cap glass tubes are preferred.
4. Transfer 10 µL of the standardised suspension into 10 mL fresh BHI (≈ $10^5$ CFU/mL). This diluted BHI suspension is the working inoculum and should be used immediately.
5. For dual-species (polymicrobial) biofilms, combine the two standardised species 1:1 at the working-inoculum step.

**Note.** Always use freshly incubated plates. Colonies left on the bench can change morphology and behaviour, and some species form a type of biofilm. In our experiments, each biological replicate was derived from different colonies on the plate.

#### 5.2 Loading the suspensions and controls

1. Dispense 160 µL of working inoculum per well in the pattern the segments will occupy. One 96-well plate accommodates a 4 × 6 layout: 4 biological replicates × up to 6 conditions.
    a. The 4 biological replicates must come from 4 independent saline suspensions.
2. Include 4 wells of catheter-only negative control (segment in sterile BHI, no bacteria).
3. Include medium-only negative controls (BHI, no catheter, no bacteria), spaced between samples, to control medium sterility and potential splatter during segment insertion.

**Note.** Keep control wells toward the far-right edge. Edge wells are marginally more likely to be brushed during washing, which can disturb biofilm integrity. Never skip these controls as they validate catheter cleaning, per plate.

#### 5.3 Confirming the inoculum by viable count

1. Serially dilute and plate the working inoculum to determine the exact starting CFU. Plate the BHI working suspension (the solution actually used).

**Note.** Do the CFU plating after the biofilm module is already in the incubator.

#### 6. Biofilm establishment and incubation

1. Remove the sterile, dried module from its bag.
2. Align the module with the freshly loaded 96-well plate and submerge the segments in the wells.
3. Using sterile tweezers or the autoclaved protective lid, press down any segment not fully seated against the well bottom.
4. Fit the 96-well plate lid on top of the module. Label only the plate or its lid, never the modular tray.
5. For microaerophilic incubation, place the plate in a 1 L gas-barrier bag with appropriate CampyGen sachet, then inside an airtight box with additional sachet. Whether microaerophilic or aerobic, use a light-tight box so the BHI is not exposed to light.
    a. These conditions approximate the restricted gas exchange and darkness of the catheterised bladder.
6. Incubate at 37°C for 24 h. Record the exact start time.
7. After 24 h, proceed to one of: (i) a further incubation round in fresh medium/condition, (ii) catheter CFU, (iii) crystal violet, or (iv) SEM. Our biofilms are considered mature after 2 × 24 h.

#### 7. Medium change and multi-day cycles

1. Prepare a fresh sterile flat-bottom 96-well plate containing 180 µL medium per well in the running layout.
2. Seat a 2 mm insert on top of the 96-well plate so that segments are lifted off the well bottom and the biofilm is not disturbed. Use 180 µL (instead of 160 µL) to compensate for the elevation.
3. Transfer the whole module onto the new plate and replace the CampyGen sachets and continue incubation.

**Note.** We printed the insert in a material which is not autoclavable, which is why we store it in isopropanol and let it air-dry briefly before use, so no isopropanol is carried into the medium or onto the segments.

When the medium surrounding the segments is itself a readout (dispersal/planktonic count), sample it at each medium change before discarding.

### 8. Readout A: viable count (CFU)

#### 8.1 Catheter-associated biofilm CFU

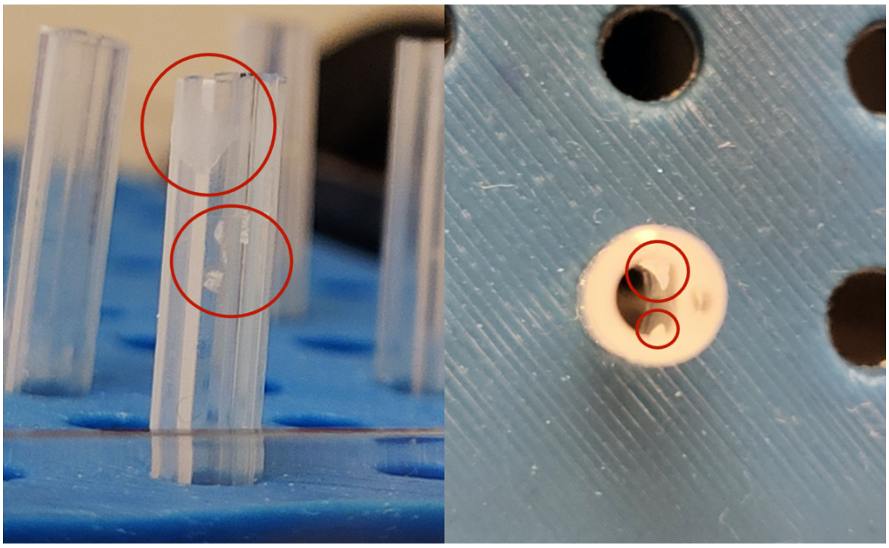

1. Place the module over a new empty sterile 96-well plate, keeping the 2 mm insert in place to protect the biofilm.
2. Wash by running 200 µL sterile PBS through the visible lumen top of each segment so it flows through the entire lumen and floods the segment above the previous medium line.
3. Empty the plate, or transfer the module to a fresh plate, and repeat, for a total of 3 × 200 µL PBS washes.

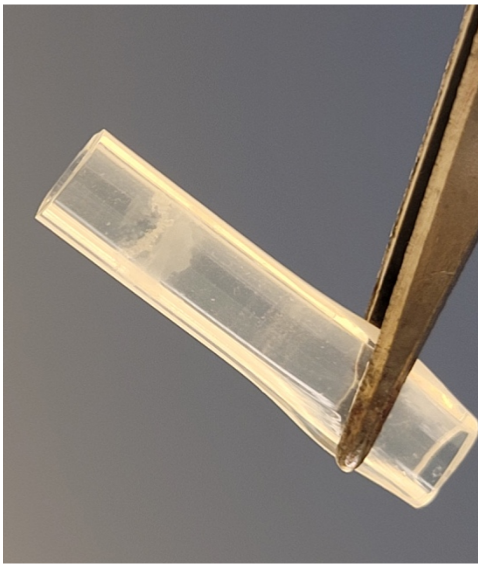

4. Prepare one 2 mL round-bottom tube per segment (e.g., 24 for a 4 × 6 layout) containing 600 µL PBS.
5. Flip the module upside down, resting on top of the 96-well lid, and unclip the tray modular part. Lift it carefully not to touch any segments.
6. With sterile tweezers, grip each segment high above the biofilm line (close to the silicon-mat) and place it in its tube. Between segments, disinfect tweezers by dipping in ethanol, flaming, and cooling in PBS.

7. Vortex the tubes 3 min at maximum speed on a multi-tube vortex.
    a. Vortexing beyond 2 min does not raise CFU yield but releases more matrix. Beyond 3 min it begins to reduce viable/culturable *E. faecalis* cells.
8. Transfer 200 µL into the A-row of a 96-well plate with rows B–H pre-filled with 180 µL PBS. Perform an 8-step tenfold serial dilution using 20 µL transfers.
9. With a multichannel pipette, spot 8 × 5 µL droplets (one per dilution) onto a pre-dried agar plate. Plate 4 technical replicates (1 plate per biological replicate). For polymicrobial samples use chromogenic Brilliance UTI agar to distinguish species.
10. Let droplets absorb (≥ 20 min) before inverting. Incubate overnight at 37°C.

**Critical.** Plates must be thoroughly dry. The 5 µL droplets sit close together, and any surface moisture makes them coalesce, making counting impossible.

#### 8.2 Surrounding-medium CFU (dispersed / planktonic cells)

Sample the medium around the segments at each medium exchange. Each sample represents only the cells released into that 24 h window.

1. Transfer 100 µL of the surrounding broth into the A-row of a 96-well plate with rows B–H pre-filled with 180 µL PBS. Perform an 8-step tenfold serial dilution using 20 µL transfers.
2. Spot 8 × 5 µL droplets per sample onto pre-dried agar (chromogenic agar for polymicrobial), 4 technical replicates. Absorb ≥ 20 min, incubate overnight at 37°C.

#### 8.3 Optical density of the surrounding medium

1. Measure OD 620 nm of the residual broth in the plate (e.g., Multiskan FC, 10 s shake before reading) to flag well-to-well deviations and potential contamination.

### 9. Readout B: crystal violet biomass

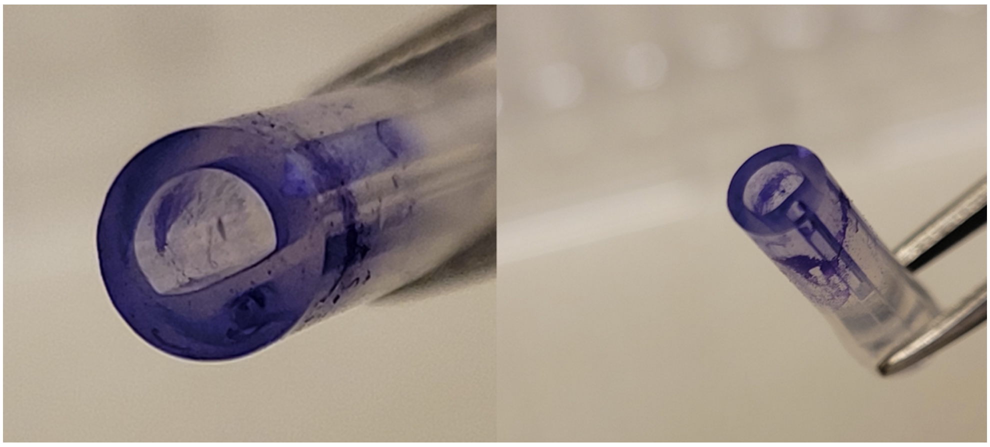

1. Place the module over a new (new but non-sterile is acceptable) 96-well plate.
2. Wash 3 × 200 µL PBS through the lumen, keeping the 2 mm insert in place. Do not rush the washes.
3. Dry the mat with segments upside down on tissue paper at 50°C until completely dry (≈ 2 h). All air bubbles must clear, as trapped air absorbs stain.
   a. A convenient way to dry after washing. Lift the tray off the 96-well plate, invert so the mat with segments drops into the lid, release and remove the tray, and let the segments dry in that position.
4. In a fume hood, prepare 0.05% crystal violet (1:20 from 1% stock in water).
5. Place individual segments into separate wells of a 96-well plate.
6. Add 200 µL crystal violet through each lumen so it flows through and surrounds the segment. Stain 15 min at room temperature.
7. Transfer stained segments to fresh wells and wash 3 × 200 µL PBS through the lumen, moving segments between empty wells until the run-through is clear and no droplet of unbound stain remains (residual stain gives false positives).

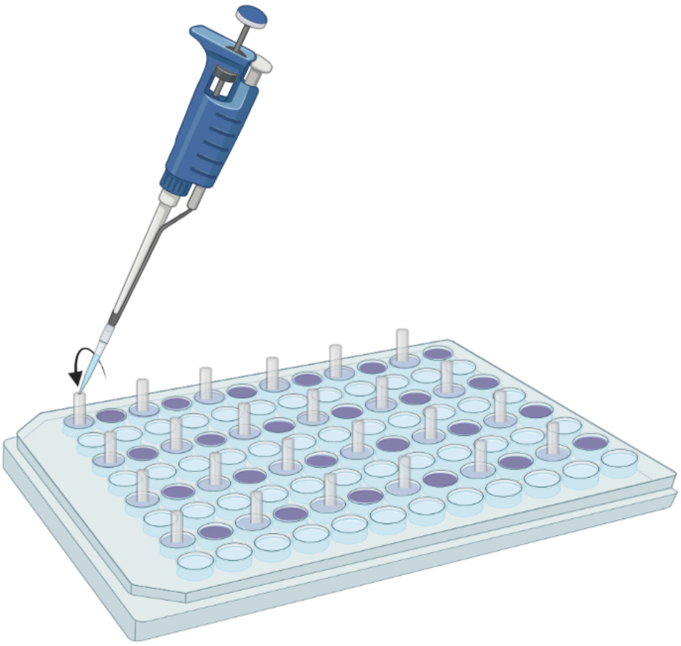

8. Blot the segment tip on paper towel to expel the last liquid (segments may be photographed here if desired).
9. Place each segment in a 2 mL round-bottom tube with 200 µL 95% ethanol and vortex 10 min to solubilise the stain. Transfer 100 µL to a microplate and read OD 540 nm.
    a. Use ethanol, not isopropanol (different refractive index in the reader).
10. Subtract the mean background (stained catheter-only controls without growth) from each sample.

**Critical.** Do not leave segments in crystal violet longer than 15 min. Staining becomes permanent and the segment must be discarded.

### 10. Readout C: scanning electron microscopy

#### 10.1 Fixation and dehydration

1. Keep the 2 mm insert in place throughout, so fixation does not damage the biofilm. Place the module over a new empty sterile 96-well plate.
2. Wash by running 220 µL sterile PBS once through the top of each segment.
3. Transfer the module to a fresh plate and wash 3 × 10 min in 250 µL phosphate buffer (Sörensen's phosphate buffer).
4. Fixate in 250 µL per segment of phosphate buffer with 2.5% glutaraldehyde + 1% paraformaldehyde for 1.5 h at room temperature. Add 200 µL to each well first, insert the segment, then add 50 µL through the lumen so it fills.
5. Dehydrate through graded ethanol, 10 min each, in a fresh plate per step: 30%, 50%, 70%, 80%, 90%, then 99.5% for 20 min.
6. Air-dry a few hours, then store in a box with desiccant (silica) beads. Image within 24 h to avoid rehydration.

#### 10.2 Mounting, coating, imaging

1. Attach conductive carbon tape to SEM pin stubs. Fix segments with tweezers. Placing two biological replicates per pin increases the chance of finding informative regions.
2. For intact (uncut) segments, an additional strip of carbon tape across the outer surface may be needed for conductivity.
3. Sputter-coat with Au/Pd (e.g., Polaron SC7640, 20 mA, 2 kV, ≈ 5–6 nm).

4. Image on a field-emission SEM (e.g., Zeiss Merlin) at 3, or 5 kV accelerating voltage. Both InLens and High-Efficiency SE2 (HE-SE2) detectors were used.

**Note.** SEM is performed with trained microscopy-facility staff. Discard any segment used for fixation.

### 11. Catheter cleaning and reuse

All segments are cleaned by this procedure regardless of prior use. Segments used for SEM fixation are discarded.

1. Place segments in a 50 mL tube of isopropanol (fully covered).
2. Vortex 10 min at maximum speed, then invert on an automatic rotator overnight.
3. Transfer segments to a fresh 50 mL tube of 5% Contrad 70 (in $dH_2O$). Discard the isopropanol to the sink.
4. Vortex 10 min at maximum speed, then invert overnight.
5. Rinse segments in distilled water (fresh 50 mL tube, vortexing) until no Contrad remains. Collect waste, do not empty down the sink.
   a. Do not reuse the Contrad 70.
6. Dry segments on a rack (e.g., an empty tip box) at 50°C overnight. Segments are then ready to remount, bag, and autoclave.

**Note.** Give newly cut segments the full wash for equal handling.
**Note.** Segments are reusable but bacterial-growth outcomes can drift after ~10 autoclave cycles. Track use per batch and discard segments after 8 uses, or earlier if dented or stained.

### 12. Experiment-specific modules

Each module plugs into the core assay at the indicated point.

#### 12.1 Conditioned (spent) medium

1. Inoculate 30 mL BHI in a 50 mL tube with a suspension standardised to 0.5 McFarland. Incubate 37°C, 24 h, shaking.
2. Centrifuge 5000 × g, 10 min, 4°C to pellet cells. Filter-sterilise the supernatant (0.22 μm).
3. Confirm sterility by plating 100 μL of the filtrate on BHI agar.

4. Use the conditioned medium in place of fresh BHI for biofilm growth (2 × 24 h), including at the 24 h medium change.

##### 12.2 Pre-conditioned surface

1. Submerge sterile catheter segments in monospecies or polymicrobial conditioned medium. Incubate 37°C, 24 h, to let soluble components coat the surface.
2. Transfer the pre-conditioned segments into fresh BHI inoculated with the test strain and grow 24 h at 37°C to assess biofilm initiation.

##### 12.3 Sequential colonisation

1. Inoculate segments with a single species and incubate 24 h for primary surface colonisation.
2. Transfer segments to fresh wells containing BHI inoculated with the second species at the standard inoculum. Incubate a further 24 h before harvesting biofilm and surrounding medium.

##### 12.4 Growth in human urine or in NaCl-diluted BHI

1. Supplement BHI with pooled human urine or 0.9% NaCl to a final 50%. Grow 24 h, transfer to fresh 50%-urine/NaCl BHI, and grow a further 24 h.

##### 12.5 Antibiotic (piperacillin–tazobactam) exposure and recovery

Model urinary TZP exposure of a mature biofilm and its subsequent recovery.

1. Prepare piperacillin (40 g/L) and tazobactam (5 g/L) stocks separately, filter-sterilised. Combine to a working solution at the clinical 8:1 ratio: 4 g/L piperacillin + 0.5 g/L tazobactam.
   a. This models the urinary concentration expected from the standard 12 g piperacillin / 1.5 g tazobactam per 24 h regimen. Here it is ≈ 1000× and ≈ 250× the BHI MICs of *E. coli* and *E. faecalis* respectively.
2. Grow a mature biofilm (2 × 24 h), then transfer the module into a plate containing the TZP working solution and incubate 24 h at 37°C.
3. Transfer the module into antibiotic-free BHI for a 24 h recovery, then repeat once for a second 24 h recovery (48 h total recovery).
4. Sample the surrounding medium after each exchange (post-exposure and after each recovery period) for dispersed-cell CFU. Harvest catheter-associated biofilm from separate parallel replicate sets at the chosen timepoints.

**Note.** Run a planktonic control in parallel: cells not derived from biofilm, exposed to the same TZP concentration in broth, to confirm that any survival is biofilm-dependent.

#### 13. Troubleshooting and critical points

- **Fused CFU droplets:** plates were not dry enough, always use pre-dried, pre-warmed plates.
- **Biofilm loss during washing:** washing too forcefully, run PBS gently through the lumen, avoid flow-back.
- **Variable crystal violet:** trapped air bubbles or unsealed inflation lumina retaining stain. Dry fully before staining.
- **Enterococcus underperforming:** light- or heat-aged BHI. Use fresh, dark-stored medium.
- **Drifting growth over time:** catheter segments past ~8 autoclave cycles. Track and discard.

#### 14. Source, attribution, and accessory files

The assay adapts the 3D-printed FlexiPeg modular system of Zaborskyte et al. (2021), modified for clinical silicone catheter segments. The original accompanying STL files (individual peg, silicone-mat mould, 96-well lid and peg-lid top) all belong to original authors and are required to reproduce the system. The 2 mm insert is an addition compatible with the original files. Variations were developed and optimised by the study authors, input on the assay was provided by G. Zaborskyte, K. Hjort and N. Kavalopoulos, and on SEM by L. Hong and H. Zhou.
